# Simultaneous infection with porcine reproductive and respiratory syndrome and influenza viruses abrogates clinical protection induced by live attenuated porcine reproductive and respiratory syndrome vaccination

**DOI:** 10.1101/2021.08.12.456007

**Authors:** Tiphany Chrun, Emmanuel A. Maze, Eleni Vatzia, Veronica Martini, Basu Paudyal, Matthew Edmans, Adam McNee, Tanuja Manjegowda, Francisco J. Salguero, Nanchaya Wanasen, Surapong Koonpaew, Simon P. Graham, Elma Tchilian

## Abstract

The porcine respiratory disease complex (PRDC) is responsible for significant economic losses in the pig industry worldwide. Porcine reproductive and respiratory syndrome virus (PRRSV) and swine influenza virus are major viral contributors to PRDC. Vaccines are cost-effective measures for controlling PRRS, however, their efficacy in the context of co-infections has been poorly investigated. In this study, we aimed to determine the effect of PRRSV-2 and swine influenza H3N2 virus co-infection on the efficacy of PRRSV modified live virus (MLV) vaccination, which is widely used in the field. Following simultaneous challenge with contemporary PRRSV-2 and H3N2 field isolates, we found that the protective effect of PRRS MLV vaccination on clinical disease and pathology was abrogated, although viral load was unaffected and antibody responses were enhanced. In contrast, co-infection in non-immunized animals reduced PRRSV-2 viremia and H3N2 virus load in the upper respiratory tract and potentiated T cell responses against both PRRSV-2 and H3N2 in the lung. Further analysis suggested that an upregulation of inhibitory cytokines gene expression in the lungs of vaccinated pigs may have influenced responses to H3N2 and PRRSV-2. These findings provide important insights into the effect of viral co-infections on PRRS vaccine efficacy that may help identity more effective vaccination strategies against PRDC in the field.

## Introduction

Porcine reproductive and respiratory syndrome (PRRS) is a viral disease responsible for major economic losses in the global pig industry (1). The disease can be subclinical depending on the strain (2, 3), however typical clinical signs are reproductive failure in sows, respiratory distress and reduction of growth performance in weaned and growing pigs (4). Mortality can be observed in infected piglets, with rates ranging from 7.5-18.5 % (5) and in some cases up to 75.5 % (6). The etiologic agents are PRRS viruses (PRRSV), single stranded positive RNA viruses from the *Arteriviridae* family (7). The first clinical description of PRRS dates to the late 1980’s, with genetically distinct PRRSV isolates described in Europe and North America, which are now recognized as two separate species PRRSV-1 (*Betaarterivirus suid 1*) and −2 (*Betaarterivirus suid 2*), respectively (7). Both species have since spread globally, but PRRSV-1 remains predominant in Europe, while PRRSV-2 predominates in the Americas and Asia. The rapid evolution of PRRSV due to a high mutation rate when replicating its genome, and recombination between strains have resulted in substantial genetic diversity, which poses challenges for the control of PRRS by vaccination (8).

Vaccination is widely practiced as one of measures used to prevent and control PRRS. Commercially available vaccines include inactivated and live attenuated (also known as modified live virus; MLV) vaccines for both PRRSV-1 and −2 (9). Although inactivated vaccines are considered safer, MLVs are preferentially used for their higher protective efficacy (8, 10). Studies have demonstrated an induction of a high level of PRRSV-specific antibodies which is associated with clinical protection against challenge infection with related PRRSV strains (9), but rather weak T cell responses after MLV immunization (11). Passive transfer of purified IgG from PRRS-convalescent pigs have suggested that vaccine-induced protection against reproductive failure and vertical transmission *in utero* is also mediated by antibodies (12, 13). Since PRRS MLV vaccine-induced antibodies often lack strong virus neutralizing properties *in vitro* (14), non-neutralizing antibody functions may also contribute but these remain poorly defined (15).

Multiple infections of pigs tend to occur naturally in the field (reviewed in (16)). Indeed, respiratory diseases in pigs in the field are often multifactorial, involving mixed infections with different viral and/or bacterial pathogens, defined as the porcine respiratory disease complex (PRDC). PRRSV and swine influenza A virus (swIAV), an enveloped single stranded segmented negative RNA virus within the *Orthomyxoviridae* family (7), are important contributors to the PRDC. Primary infection with PRRSV and swIAV leads to pneumonia caused by opportunistic pathogens, such as *Pasteurella multocida, Mycoplasma hyopneumonia*, *Actinobacillus pleuropneumoniae* and *Bordetella bronchiseptica* (16). Previous *in vivo* PRRSV/swIAV co-infection or superinfections studies demonstrated a potentiation of disease compared to single infection (17, 18). It was recently shown that concomitant swIAV infection modulated the immune response to PRRS MLV vaccination, albeit without impacting efficacy (19), but it remains unknown whether such viral co-infection interferes with protection conferred by PRRS vaccination.

In the present study, we examined the efficacy of PRRS MLV vaccination to PRRSV-2/H3N2 co-infection. We evaluated clinical signs, viral load, PRRSV-2-specific antibody and T cell responses in PRRS MLV-vaccinated pigs challenged simultaneously with contemporary field-isolated PRRSV-2 and swIAV H3N2 strains. We report here that PRRSV-2/H3N2 co-infection abrogated the protective effect of PRRS MLV vaccine on lung pathology, although it did not alter viral load. Moreover, PRRS MLV reversed the beneficial effect of PRRSV-2/H3N2 co-infection in decreasing H3N2 lung viral loads, highlighting a potential interference of PRRS MLV vaccination on the subsequent host immune response against H3N2.

## Materials and methods

### Cell lines

Madin–Darby canine kidney (MDCK) cells were cultured in Eagle’s minimum essential medium (MEM, Merck, Feltham, UK) supplemented with 10 % heat-inactivated fetal bovine serum (HI FBS, Thermo Fisher Scientific, Loughborough, UK) and antibiotics (100 U/mL penicillin and 100 µg/mL streptomycin, Thermo Fisher) at 37 °C in a humidified 5 % CO_2_. African green monkey kidney (MARC-145) cells were cultured in Dulbecco’s modified MEM (DMEM, Merck) supplemented with 10 % HI FBS and antibiotics at 37 °C in a humidified 5 % CO_2_ atmosphere.

### Vaccine and virus strains

PRRSV-2 16CB02 was isolated from pig serum collected from a diseased farm in Chonburi, Thailand in 2016. The serum was adsorbed onto MARC-145 cells and cultured in OptiMEM (Thermo Fisher Scientific) supplemented with 10 % FBS. After 72 h, cytopathic effect (CPE) was observed, and the supernatant was collected. Cells were then infected with a diluted culture supernatant before being overlayed with MEM supplemented with 0.3 % bovine serum albumin (BSA, Merck), 0.22 % sodium bicarbonate (Merck), 1 % penicillin/streptomycin (Merck) and 1 % methylcellulose (Merck). The PRRSV-2 16CB02 isolate (referred to as PRRSV-2) was derived from a single plaque-derived culture, which resulted in obvious CPE after 96 h incubation. Partial genome (ORFs 2-7) sequencing was performed (GenBank, accession number MZ700336) and the ORF5 sequence was compared with sequences in the NCBI database using BLAST showed that 16CB02 belonged to lineage 8.7 of PRRSV-2 (*Betaarterivirus suid 2*). PRRSV-2 16CB02 was propagated in MARC-145 cells by infection with a multiplicity of infection (MOI) of 0.08 for 1 h at 37 °C in 5 % CO_2_. After removing the inoculum cells were incubated for 3 days in DMEM with 10 % FBS and antibiotics at 37 °C in 5 % CO_2_. Supernatant was harvested, centrifuged at 500 × g for 10 min and stored at −80 °C until use. The viral stock was titrated and expressed as TCID_50_/mL.

SwIAV H3N2 CM5 strain was isolated from a pig farm in the Province of Lumpoon, Thailand in 2018. Nasal swab samples were first screened for IAV by one-step RT-PCR. Viral RNA was extracted from samples with the GenUP™ Virus RNA kit (biotechrabbit, Henningsdorf, Germany). One-step RT-PCR was conducted using matrix (M) segment specific primers (20) and performed on a T100™ Thermal Cycler (Bio-Rad) utilizing the PrimeScript™ One-Step RT-PCR kit (Takara Bio, Shiga, Japan). PCR products were then separated and visualized by agarose gel electrophoresis. The RT-PCR-positive nasal swab samples were then subjected to IAV isolation using MDCK cells. CPE was observed daily, and CPE positive supernatants collected and confirmed by HA test and RT-PCR for M gene detection as previously described (21). Sequencing of H3N2 RNA segments were performed and alignment of HA (GenBank, accession number MZ665044) and NA (GenBank, accession number MZ665046) sequences against sequences in the NCBI database revealed that CM5 strain was a H3N2 subtype. H3N2 CM5 (referred to as H3N2) was propagated by infecting MDCK cells at MOI 0.001 in serum-free MEM and incubated at 37 °C in 5 % CO_2_ for 1 h. After washing the cell monolayer, cells were incubated in serum-free MEM supplemented with 2 µg/mL TPCK-treated trypsin (Merck) for 2 days at 37 °C in 5 % CO_2_. Supernatant was harvested, centrifuged at 880 × *g* for 10 min and stored at −80 °C until use. The viral stock was titrated by plaque assay and expressed as pfu/mL.

PRRS MLV (Ingelvac® PRRS MLV, Boehringer Ingelheim Vetmedica GmbH, Ingelheim am Rhein, Germany) was used following reconstitution according to manufacturer’s instructions.

### Immunization and challenge study

The animal experiment was approved by the Animal Welfare and Ethical Review Body at The Pirbright Institute, UK. The treatment, housing, husbandry, and procedures were performed in accordance with the UK Animal (Scientific Procedures) Act 1986 (Project Licence P6F09D691). Thirty-six, 5-7 weeks-old, Large White-Landrace-Hampshire crossbred female pigs were sourced from a high health status commercial herd and were housed in a high biocontainment facility at The Pirbright Institute. Animals were tested for the absence of exposure to IAV and PRRSV prior to their arrival by serological tests via hemagglutination inhibition test against four standard IAV antigens (pdmH1N1, H1N2, H3N2 and avian-like H1N1 strains), and antibody (Ab) ELISA and RT-PCR tests against PRRSV (Animal and Plant Health Agency, Weybridge, UK). Pigs were randomly assigned to 6 groups of 6 pigs each, which were untreated (naïve), immunized with 2 mL containing 10^4.5^ TCID_50_ of Ingelvac® PRRS MLV (Vac) or with 2 mL of PBS (Ctrl) by intramuscular (i.m.) injection (**Figure 1A**). On day 33 post-vaccination (dpv), Ctrl and Vac pigs were inoculated intranasally with 4 mL (2 mL/nostril) containing 5 × 10^6^ pfu of swIAV H3N2 CM5 (Ctrl + H3N2), 10^5^ TCID_50_ PRRSV-2 16CB02 (Ctrl + PRRSV-2 or Vac + PRRSV-2), or concurrently with 5 × 10^6^ pfu swIAV and 10^5^ TCID_50_ PRRSV-2 (Ctrl + PRRSV-2/H3N2 or Vac + PRRSV-2/swIAV) diluted in DMEM using a mucosal atomization device (MAD 300, Wolfe Tory Medical, Salt Lake City, USA). To analyze the immune response, pigs from Ctrl + PRRSV-2, Vac + PRRSV-2 and Vac + PRRSV-2/H3N2 groups were bled at 0 dpv, then pigs from all groups at 20 and 38 dpv for peripheral blood mononuclear cell (PBMC) and serum isolation. After the challenge, pigs were observed twice per day until the end of the study for monitoring and scoring of clinical signs (**Supplementary Table S1**). Rectal temperatures were taken on −1, 0, 1, 2, 3 and 4 days post-challenge (dpc). Nasal swabs were collected daily for viral detection. At 5 dpc, the animals were humanely euthanized with an overdose of pentobarbital sodium anesthetic.

**Figure 1:**
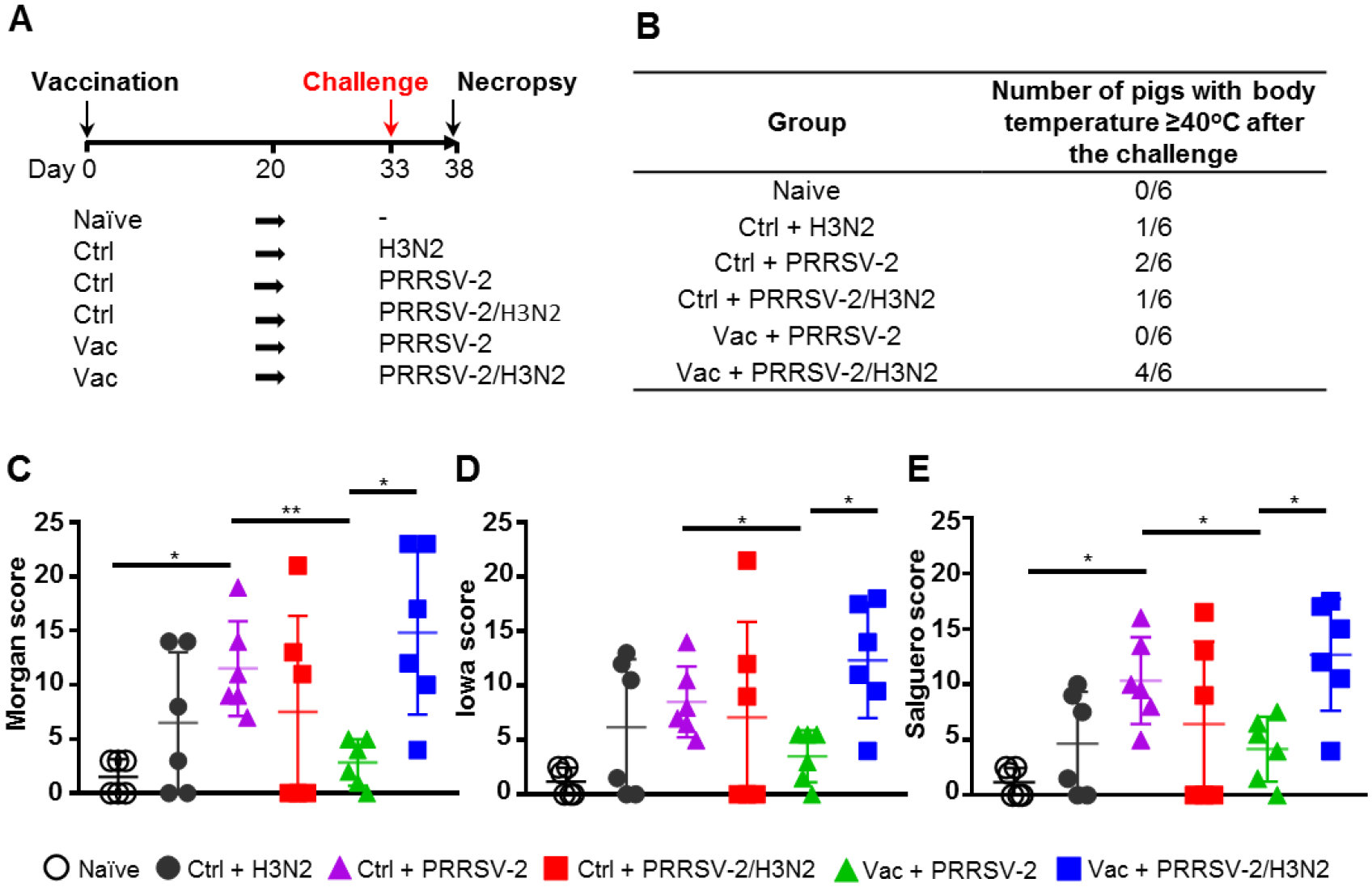
Clinical signs and lung lesions. **(A)** Pigs were vaccinated intramuscularly with Ingelvac PRRS^®^ MLV (Vac) or with PBS (Ctrl) or were untreated (naïve group). Thirty-three days after the vaccination, pigs were challenged by intranasal inoculation with H3N2, PRRSV-2 or simultaneously with PRRSV-2/H3N2. Nasal swabs were daily taken after the challenge and pigs were culled 5 days later (38 days post-vaccination (dpv)). Sera and PBMC were collected at 0, 20 and 38 dpv. Clinical signs and rectal temperature were monitored daily after the challenge (dpc). **(B)** Table indicating the number of pigs that developed fever after the challenge. **(C-E)** Lungs sections were scored for histopathological lesions (**C**; Morgan score), lesions with presence of influenza NP-positive cells (**D**; Iowa score) or lesions with presence of PRRSV N-positive cells (**E**; Salguero score). Each symbol represents an individual animal within the indicated group (n=6 per group). The horizontal lines represent mean ± SD. Comparisons between 2 group were analyzed using Mann-Whitney test and asterisks indicate significant differences (*p < 0.05; **p < 0.01).

**Table 1:**
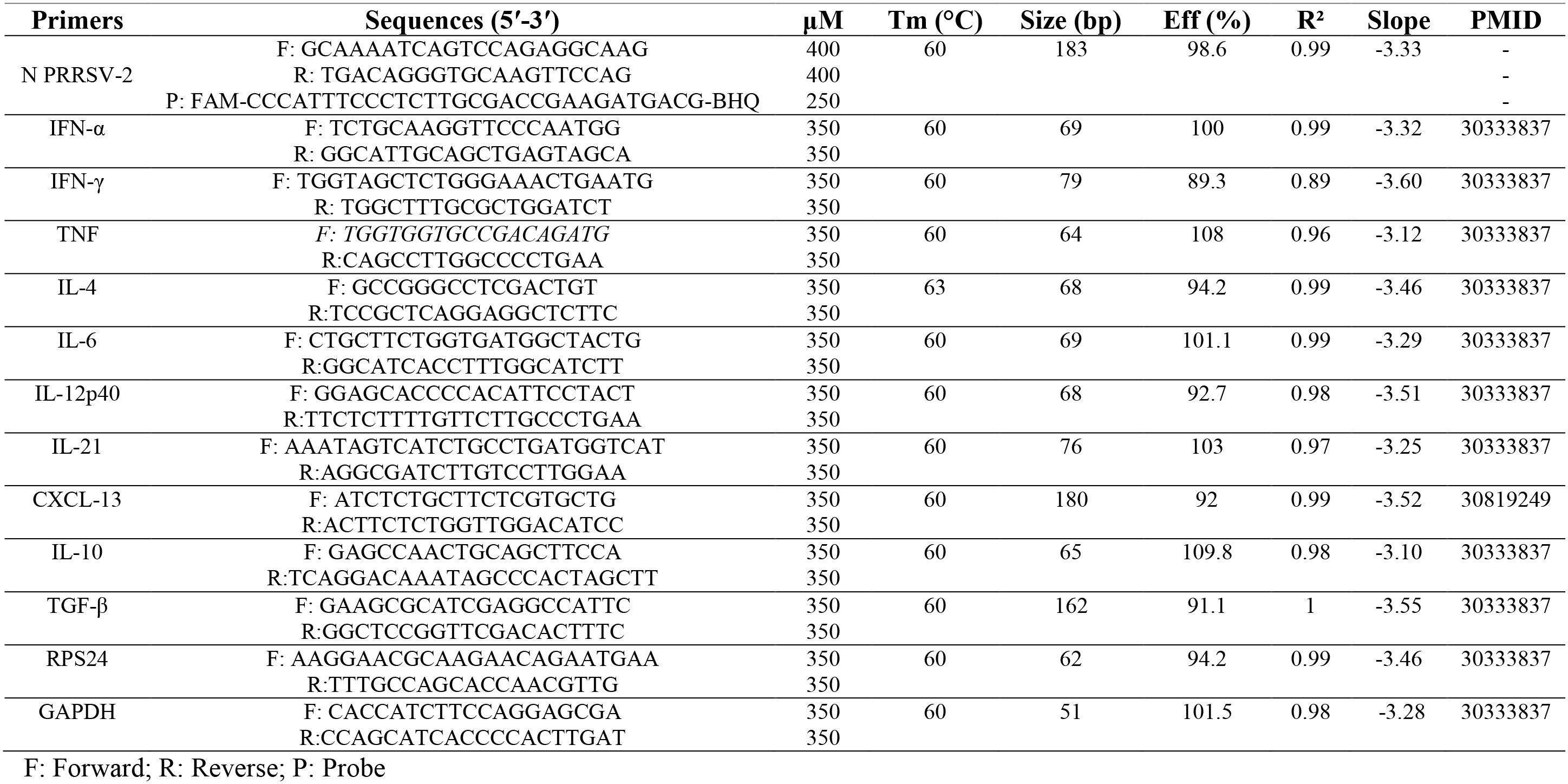
Primers used for RT-qPCR and RT-PCR

### Pathological and histopathological examination of lungs

Lungs were removed post-mortem and tissue samples were taken from cranial, cardiac, and diaphragmatic lobes of the left lung, and immersed in 10 % neutral buffered formalin for fixation and histological processing. Formalin fixed tissues were paraffin wax-embedded and 4µm sections cut and stained with hematoxylin and eosin (H&E). Immunohistochemical staining of IAV nucleoprotein (NP) and PRRSV nucleoprotein (N) protein was performed in 4 µm tissue sections as previously described (22, 23). Histopathological changes in stained lung tissue sections were scored by a board-certified veterinary pathologist blinded to the treatment group. Lung histopathology was scored (“Morgan score”) using five parameters (necrosis of the bronchiolar epithelium, airway inflammation, perivascular/bronchiolar cuffing, alveolar exudates, and septal inflammation) scored on a 5-point scale of 0 to 4 and then summed to give a total slide score ranging from 0 to 20 and a total lung score from 0 to 60 (24). Two additional scoring systems were used to analyze the contribution of IAV (“Iowa”) and PRRSV-2 (“Salguero”) in the lung lesions. Abundance of viral antigen was assessed using the influenza NP staining (HB-65 mAb, Bio X Cell, Lebanon, USA) and was scored as described previously (25). Similarly, PRRSV N protein staining was performed (SDOW17-A mAb, Rural Technologies, Brookings, USA) and the scoring method was adapted from the Iowa IAV scoring method as described above.

### Sample collection and cell isolation

Blood samples were collected at 0, 20 and 38 dpv using BD Vacutainer™ SST™ II Advance Tubes and BD Vacutainer Heparin Blood Collection Tubes (both Fisher Scientific). Serum tubes were centrifuged at 880 × *g* for 10 min and the resulting serum was aliquoted and stored at −80 °C. Heparinized blood diluted 1:1 in PBS overlaid onto Lymphopure density gradient medium 1.077 g/mL (BioLegend, San Diego, USA) in Leucosep tubes (Greiner Bio-One, Gloucestershire, UK) and centrifuged at 800 × *g* for 15 min. PBMC were harvested from the interface and washed with PBS. Erythrocytes were removed with RBC Lysis Buffer (BioLegend). The cell suspension was washed again, filtered through a 100 µm cell strainer and cryopreserved in freezing medium (HI FBS with 10 % DMSO). Bronchoalveolar lavage (BAL) was performed on the isolated right lung with 300 mL of PBS. Cells were isolated by centrifugation of the BAL fluid (BALF) at 490 × *g* for 5 min, filtered through 100 µm cell strainer, and cryopreserved in freeing medium. The BALF supernatant was aliquoted, and frozen at −80 °C for virus titration. For gene expression analysis, pieces of cranial lung lobe tissues were submerged into RNAlater™ Stabilization Solution (Thermo Fisher Scientific), placed at room temperature for 6 h and then stored at −80 °C until use. Sampling of nasal secretions was performed at 0, 1, 2, 3, 4 and 5 dpc using cotton swabs (one in each nostril, Scientific Laboratory Supplies, Nottingham, UK). Swabs were then placed into virus transport medium and processed as described previously (26).

### RNA extraction from fluids and tissues

Total RNA was extracted from nasal swabs, BALF and serum using QIAamp Viral RNA kit (QIAGEN, Manchester, UK) according to the manufacturer’s instructions. Piece of lungs (200-300 mg) were weighed and homogenized using gentleMACS Octo dissociator (Miltenyi Biotec) in RPMI 1640 medium in M tubes (Miltenyi Biotec, Woking, UK). After centrifugation at 880 × *g* for 5 min, supernatants were collected. Total RNA was extracted with RNAeasy kit (QIAGEN) according to the manufacturer’s instructions. Extracted RNA were stored at −80 °C until use.

### Virus titration by plaque assay

SwIAV titers in nasal swabs and BALF were determined by plaque assay as previously described (22). MDCK cells were inoculated with 10-fold serially diluted samples in MEM. After 1 h incubation at 37 °C, cells were washed and incubated for further 72 h at 37 °C in 5 % CO_2_ under an overlay medium consisting of MEM with 0.21 % BSA, 2 mM L-glutamine, 0.15 % sodium bicarbonate, 10 mM HEPES (Merck), 1 % penicillin/streptomycin, 0.01 % Dextran DEAE (Merck), 0.6 % agar (Merck) and 2 µg/mL TPCK trypsin (Merck). The pfu were counted following staining with 1 % (w/v) crystal violet (Merck).

### Viral RNA quantification by qRT-PCR

PRRSV-2 RNA quantification in nasal swabs, BALF and serum was performed by quantitative reverse-transcriptase PCR (qRT-PCR) using the one-step Quantinova™ Probe RT-PCR kit (QIAGEN). Primers and a TaqMan probe were designed to hybridize to a sequence in ORF7 (encoding N), and sequences are shown in **Table 1**. To determine the viral RNA copies/mL in a sample, an RNA standard was generated. PRRSV-2 RNA was used to amplify cDNA encoding the full-length of N gene by RT-PCR (One-step RT-PCR, QIAGEN) using specific primers with the forward containing a T7 promoter sequence at the 5’ end. The PCR product was gel purified (Illustra™ GFX PCR DNA and Gel Band Purification Kit, Merck) and used as template for *in vitro* transcription using the MEGAscript T7 Transcription Kit (Thermo Fisher Scientific), according to the manufacturer’s protocol. After DNase treatment, the concentration of RNA was measured using a Nanodrop spectrophotometer (Thermo Fisher Scientific), and the number of RNA molecules/μL was calculated using Avogadro’s number (6.023 × 10^23^). The qRT-PCR was performed with 5 µL of the eluted samples and 15 µL of the master mix. Samples and standards were run in duplicate under the following conditions on a Stratagene Mx3500P cycler (Agilent): reverse transcription at 45 °C for 20 min, denaturation at 95 °C for 5 min, 35 amplification cycles of denaturation at 95 °C for 5 s and combined annealing/extension at 60 °C for 30 s. The viral genome copy numbers were determined by interpolation of the standard curve and limit of quantification was estimated at 60 copies/mL.

### Gene expression by RT-PCR

Total RNA samples were treated with DNA-free Kit (Thermo Fisher Scientific). Absence of residual DNA was verified by using RNA samples as a template for SYBR Green PCR. Relative mRNA expression was evaluated by RT-PCR using the QuantiNova SYBR Green RT-PCR Kit (QIAGEN). Primers of selected cytokines and chemokines used in previous studies (27, 28) are listed in **Table 1**. RT-PCR assays were validated and displayed an efficiency between 80 % and 110 %. Reactions utilized 5 µL containing 100 ng of RNA sample with 15 µL of the master mix. Samples were run in duplicate under the following conditions on a Stratagene Mx3500P cycler (Agilent): reverse transcription at 50 °C for 30 min, denaturation at 95 °C for 5 min, 40 amplification cycles of denaturation at 95 °C for 10 s and appropriate combined annealing/extension temperature shown in Table 1 for 60 s. Melting curves were generated after each run to confirm the RT-PCR assay specificity. Fold changes in gene expression were calculated using the delta-Cq method with multiple housekeeping genes (29). The housekeeping genes *RPS24* (ribosomal protein S24) and *GAPDH* (glyceraldehyde 3-phosphate dehydrogenase) were found to be stably expressed in pig lung cells (27, 30) and were used as reference genes to normalize the data. Samples from naïve group were used as calibrators.

### Intracellular cytokine staining

Cryopreserved PBMC and BAL cells were thawed, washed, and resuspended in RPMI 1640 medium with 10 % HI FBS and antibiotics. To assess the intracellular cytokine production, 2 × 10^6^ cells/well were seeded in a 96-well-round bottom tissue culture plate with complete RPMI 1640 medium. After 5 h of resting at 37 °C, cells were re-stimulated with H3N2 CM5 (MOI 0.1) and PRRSV-2 16CB02 (MOI 0.1) for 18 h at 37 °C. Cells cultured in complete RPMI 1640 medium only served as negative controls. BD GolgiPlug at 1:1,000 (BD Biosciences, Wokingham, UK) was added into the well for a further 4 h before staining. Cells stimulated with a cocktail of PMA (phorbol 12-myristate 13-acetate) (20 ng; Merck), ionomycin (500 ng; Merck) and BD GolgiPlug at 1:1,000 for 4 h was used as positive a control. Cells were washed and stained with 50 µL of surface marker antibodies listed in **Supplementary Table S2**, and Near-IR Fixable LIVE/DEAD stain (Thermo Fisher Scientific) diluted in PBS with 2 % FBS for 20 min at 4 °C. After two washing steps, the cells were fixed and permeabilized (BD Cytofix/Cytoperm kit; BD Biosciences) for 30 min at 4 °C. Intracellular cytokine staining (ICS) was performed according to the manufacturer’s directions using 50 µL of cytokine mAbs listed in **Supplementary Table S2** diluted in Perm/Wash Buffer (BD Biosciences) for 30 min at 4 °C. Subsequently, cells were washed in Perm/Wash Buffer and fixed with 2 % PFA in PBS (Santa Cruz Biotechnology, Heidelberg, Germany). Cells were analyzed with a BD LSRFortessa Flow Cytometer (BD Biosciences). Analysis was performed with FlowJo version 10.6.2 (FlowJo, LLC). Compensation was set according to single color staining controls. Isotype controls and fluorescence minus one (FMO) controls were used to validate the staining and to set the gates.

### Evaluation of anti-PRRSV antibody responses by ELISA and virus neutralization assay

To detect anti-PRRSV IgG, HI sera were tested using PrioCHECK PRRSV Antibody ELISA Kit (Thermo Fisher Scientific) for detection of PRRSV N protein-specific antibodies, according to the manufacturer’s instructions. Optical density (OD) was measured at 450 nm using a microplate reader (Synergy™ HT Multi-Detection Microplate Reader, BioTek Instruments, USA). Percentage positivity (PP) was calculated using following formula: PP = (OD of sample – OD of negative control) / (OD of positive control – OD of negative control) × 100. The cut-off was determined according to the supplier’s protocol i.e., samples above 30 PP were considered as positive. To measure anti-PRRSV IgG titers in serum, an ELISA using PRRSV-2 infected cell lysate as antigen was used. Antigen was generated as described elsewhere (31). Briefly, lysate was obtained by sonicating PRRSV-2 16CB02 infected MARC-145 cell pellets in lysis buffer (1 % Triton X-100, 50 mM borate, 150 mM NaCl, pH 9). The supernatant was clarified and stored at −80 °C. High-binding 96-well plates (Nunc Maxisorp^TM^, Thermo Fisher Scientific) were coated overnight at 4 °C with 100 µL of lysate diluted at 1:40 in carbonate buffer (Merck). After a blocking step with PBS with 4 % milk for 1 h, serial 2-fold dilutions of HI serum were added in duplicates, starting from 1:40 and incubated for 1 h at room temperature (RT). Bound PRRSV-2-specific IgG was detected using goat anti-pig IgG conjugated with horseradish peroxidase (HRP; Bio-Rad) diluted at 1:10,000 in blocking buffer for 1 h at RT. HRP enzymatic activity was revealed using 3,3’,5,5’-tetramethylbenzidine (TMB) substrate solution (Thermo Fisher Scientific) for 5 min and was stopped by adding 1.2 M sulfuric acid. Antibody endpoint titers were determined as the highest dilution giving twice the OD of the negative control wells (lysate coated well only). PRRSV neutralizing Ab titers were assessed using an adapted protocol previously described (32). Briefly, serial 2-fold dilutions of HI serum incubated with 400 TCID_50_ of PRRSV-2 for 1 h at 37 °C were added to MARC-145 cell monolayers. After 3 days incubation at 37 °C, cells were fixed and permeabilized (2 % PFA 0.1 % Triton X-100 in PBS) for 10 min at RT and blocked with 10 % goat serum in PBS. Cells were stained using an anti-PRRSV N mAb (SDOW17-A, Rural Technologies) diluted 1:1,600, followed by a secondary goat anti-mouse IgG conjugated to HRP (Bio-Rad) diluted 1:1,000. PRRSV-2 positive cells were revealed using 3, 3’-diaminobenzidine substrate (DAB, Vector Laboratories, Burlingame, USA) for 10 min. Neutralizing Ab titers were calculated as the reciprocal serum dilution that neutralized viral infection in 100 % of the wells.

### Statistical analysis

Data were analyzed with GraphPad Prism 8.0.1 software. As distribution was not normal (Anderson-Darling test), the non-parametric unpaired Kruskal-Wallis test followed by Dunn’s correction was applied for multiple comparison (lung lesions scores, viral load, proportion of immune cell subsets and genes expression). The unpaired non-parametric Mann-Whitney test was used to compare data between 2 groups. The matched paired non-parametric Wilcoxon test was used to compare the T cell and antibody responses at day 0 and different timepoints within the same group.

## Results

### PRRSV-2/H3N2 co-infection abrogates PRRS MLV-induced clinical protection

To investigate whether concurrent infection with H3N2 and PRRSV-2 can interfere with the efficacy of PRRS MLV vaccination, groups of 6 pigs were immunized with a commercial PRRS MLV (Vac) or mock vaccinated with PBS (Ctrl) by intramuscular injection (**Figure 1A**). Animals were challenged at 33 day post-vaccination (dpv) with 5 × 10^6^ pfu of a swIAV H3N2 field isolate (Ctrl + H3N2), 10^5^ TCID_50_ of a PRRSV-2 field isolate (Ctrl + PRRSV-2 and Vac + PRRSV-2) or both viruses (Ctrl + PRRSV-2/H3N2 and Vac + PRRSV-2/H3N2) by intranasal inoculation. Pigs were culled 5 days post-challenge (38 dpv). A group of 6 pigs remained untreated throughout the study and served as naïve controls. Clinical signs and rectal temperature were monitored daily post-challenge (**Supplementary Tables S1 and S3**). In the unvaccinated and H3N2 infected group (Ctrl + H3N2), 1/6 pigs had an elevated temperature (≥40 °C) after the challenge, 2/6 pigs in the unvaccinated and PRRSV-2 infected group (Ctrl + PRRSV-2) and 1/6 pigs in the unvaccinated and co-infected group (Ctrl + PRRSV-2/H3N2) (**Figure 1B**). No other apparent clinical signs were observed after challenge (data not shown), which indicated that both field strain viruses induced a mild disease during this stage of infection, and that co-infection did not enhance clinical disease. None of the vaccinated pigs infected with PRRSV-2 (Vac + PRRSV-2) showed hyperthermia but 4/6 of the pigs vaccinated and co-infected with PRRSV-2/H3N2 (Vac + PRRSV-2/H3N2) developed fever (≥40 °C) which lasted for 3 consecutive days in one pig (**Supplementary Table S3**).

Following the experimental infection, microscopic examination of the lungs collected at 5 dpc revealed mild to moderate bronchointerstitial pneumonia in H3N2 (Ctrl + H3N2 or Ctrl + PRRSV-2/H3N2) or PRRSV-2 infected groups (Ctrl + PRRSV-2 or Vac + PRRSV-2/H3N2) in comparison to the naïve controls or vaccinated group and infected with PRRSV-2 (Vac + PRRSV-2) (**Supplementary Figures S1**). Lesions were characterized by bronchial epithelial necrosis, an expansion and thickening of alveolar septa, and lymphocyte cuffing and histiocytic cellular infiltration in the peribronchial and perivascular space as previously described after H3N2 (33, 34) and PRRSV (23, 35) infection. Immunohistochemistry analysis indicated that IAV NP was observed in the relevant group, i.e., mainly observed in cells within the epithelium of bronchi and bronchioles in Ctrl + H3N2, Ctrl + PRRSV-2/H3N2 and Vac + PRRSV-2/H3N2 groups, and not in naïve and Ctrl + PRRSV-2 and Vac + PRRSV-2 groups (**Supplementary Figures S2A**). Similarly, PRRSV N was localized, as expected, in alveolar and interstitial macrophages in the Ctrl + PRRSV-2, Ctrl + PRRSV-2/H3N2, Vac + PRRSV-2 and Vac + PRRSV-2/H3N2 groups (**Supplementary Figures S2B**).

The severity of lung lesions was compared between unvaccinated groups after a single and co-infection using three different scoring systems, with similar results (**Figures 1C-E**). Single infection of unvaccinated pigs with H3N2 or PRRSV-2 induced lung lesions (mean scores of 5.6 and 9.6, respectively) in comparison to naïve pigs (mean scores of 1), although significant differences in lung lesions were found only after PRRSV-2 infection (p<0.05 vs naïve). Simultaneous PRRSV-2/H3N2 infection did not significantly increase the lung lesions in the unvaccinated group (mean score of 7) compared to single virus infection groups. To assess whether PRRS vaccine conferred protection, lung lesion scores were compared between Ctrl + PRRSV-2 and Vac + PRRSV-2 groups. A significant reduction of the score was observed in Vac + PRRSV-2 (mean score of 3.5) in comparison to Ctrl + PRRSV-2 group (mean score 9.6; p<0.05 all scoring systems). However, PRRSV-2/H3N2 co-infection abrogated the protective effect of the PRRS MLV since the Vac + PRRSV-2/H3N2 animals exhibited high lung lesion scores in comparison to Vac + PRRSV-2 animals (mean score 13.2 versus 3.5, respectively) (p < 0.05, all scoring systems).

Collectively, these results indicate that co-infection with H3N2 abrogates the PRRS MLV mediated protective effect, by enhancing lung lesions and clinical disease.

### Assessment of PRRSV-2 and H3N2 viral loads

To assess whether PRRSV-2/H3N2 co-infection affects the ability of PRRS MLV vaccine to reduce virus loads, PRRSV-2 RNA was quantified in nasal swabs, BALF and serum (**Figure 2**). Shedding of PRRSV-2 was observed in nasal swabs from 3 to 5 dpc in all PRRSV-2 challenged groups and not in the samples from naïve or Ctrl + H3N2 group (**Figure 2A**). At these early timepoints post-infection, we found great variability within group. PRRSV-2 genome was detected in nasal swabs in 4/6 pigs from Ctrl + PRRSV, in 1/6 pigs from Ctrl + PRRSV-2/H3N2, in 5/6 pigs from Vac + PRRSV and in 1/6 pigs from Vac + PRRSV-2/H3N2. These data indicate that PRRS MLV did not reduce viral shedding. Of note, PRRSV-2 RNA was detected in only 1/6 in co-infected groups (Ctrl or Vac), suggesting that co-infection may reduce PRRSV-2 viral shedding.

**Figure 2:**
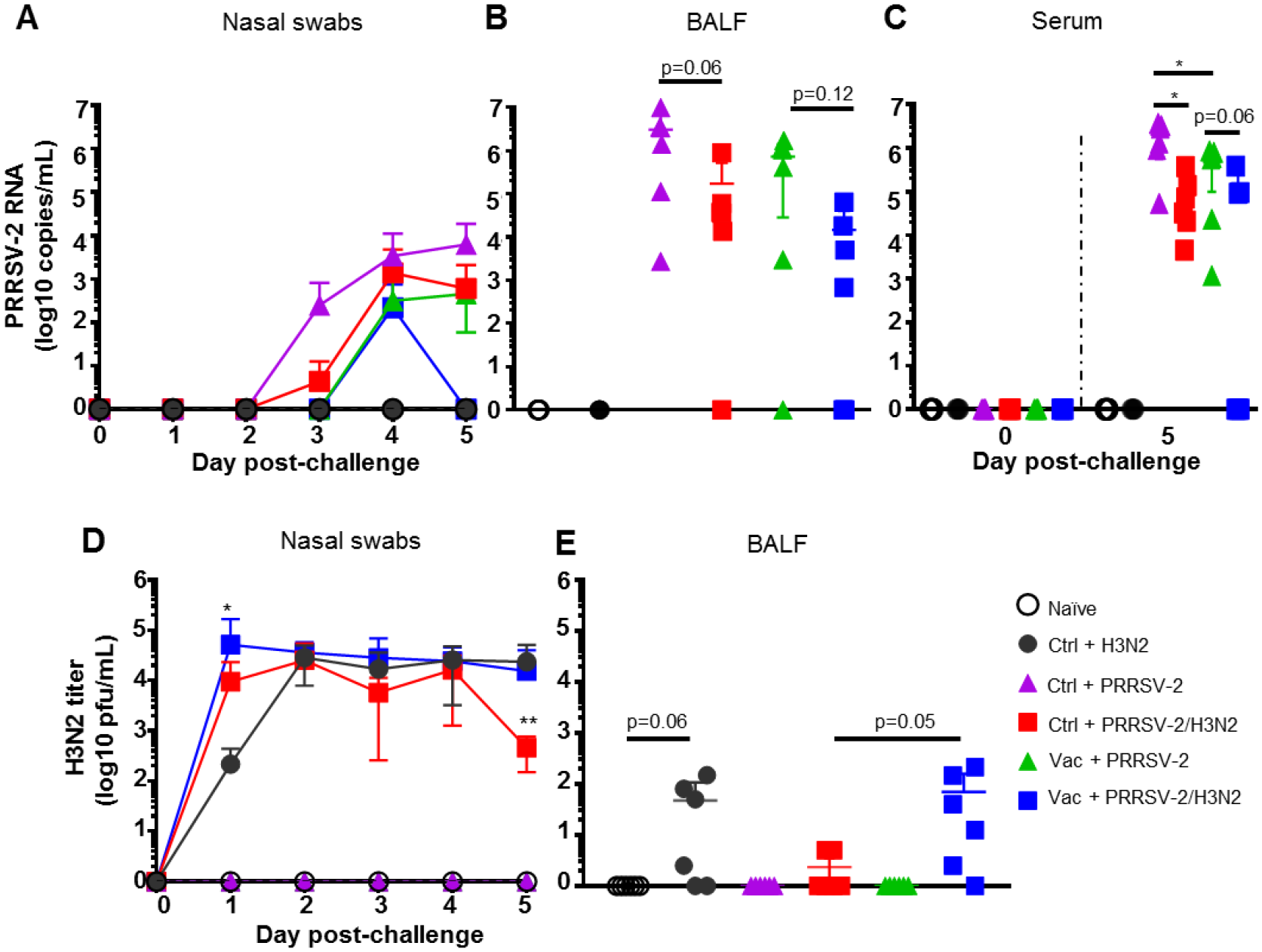
PRRSV-2 and H3N2 loads. Quantification of PRRSV-2 viral RNA in the nasal swabs (**A**), BALF (**B**) and serum (**C**) were determined by qRT-PCR. H3N2 titers in nasal swabs (**D**) and BALF **(E**) were determined by plaque assay. The mean values (**A** and **D**) or individual values (**B, C, D** and **E**) for each group ± SD are indicated (n=6 per group). P values were determined using Mann-Whitney test and asterisks indicate significant differences (*p< 0.05).

In the lower respiratory tract, high levels of PRRSV-2 RNA were measured in BALF after single PRRSV-2 infection in both Ctrl + PRRSV-2 or Vac + PRRSV-2 groups and not in naïve and Ctrl + H3N2 groups (**Figure 2B**). A reduction (average 4-fold) of the PRRSV-2 RNA load was observed in the Vac + PRRSV-2 group compared to Ctrl + PRRSV-2 group although the difference was not significant, indicating that intramuscularly administered PRRS MLV may help to limit the viral replication in the lung at 5 dpc. Interestingly, in both vaccinated and unvaccinated groups, there was a trend towards lower PRRSV-2 RNA loads in BALF following the co-infection compared to the single infection. These results indicate that co-infection seems to be efficient to reduce viral load (**Figure 2B**). In the serum, PRRSV-2 RNA was also detected at 5 dpc in Ctrl or Vac + PRRSV-2 groups and not in naïve and Ctrl + H3N2 groups (**Figure 2C**). However, levels of PRRSV-2 RNA were significantly reduced in Vac + PRRSV-2 compared to Ctrl + PRRSV-2 (mean 4-fold, p<0.05) indicating that the vaccine conferred a degree of protection. A significant reduction of PRRSV-2 viral RNA was also measured in Ctrl + PRRSV-2/H3N2 group compared to Ctrl + PRRSV-2 (mean 19-fold; p<0.05). The co-infection also slightly reduced PRRSV-2 RNA in Vac + PRRSV-2/swIAV group as compared to the Vac + PRRSV-2 group, although the difference was not statistically significant (mean 5-fold; p=0.06).

H3N2 load was also assessed in nasal swabs and BALF. Virus titers were detectable in nasal swabs of groups challenged with H3N2 (alone or with PRRSV-2) and not in naïve and PRRSV-2 only infected animals (**Figure 2D**). After a single infection with H3N2 (Ctrl + H3N2), a peak of the virus shedding was reached at 2 dpc, followed by a plateau until 5 dpc. After PRRSV-2/H3N2 co-infection, the peak of nasal shedding was also reached at 2 dpc in the Ctrl + PRRSV-2/H3N2 group, whereas this was seen earlier at 1 dpc in Vac + PRRSV-2/H3N2 group. A significant reduction of H3N2 was measured in the Ctrl + PRRSV-2/H3N2 at 5 dpc (mean 50-fold; p<0.01), however, this reduction disappeared in animals vaccinated with PRRS MLV (Vac + PRRSV-2/H3N2 group) which exhibited similar pattern of H3N2 shedding to Ctrl + H3N2.

In BALF, H3N2 was detected at low levels after the single (Ctrl + H3N2 – mean of 47 pfu/mL) or after co-infection (Ctrl + PRRSV-2/H3N2 – mean of 2 pfu/mL) although these differences were not significant (**Figure 2E**). After co-infection, PRRS MLV vaccination markedly increased the H3N2 titers in the Vac + PRRSV-2/H3N2 group group compared to the non-immunized co-infected group (mean 69 and 2 pfu/mL respectively, p=0.05).

Together, these data indicated that PRRS MLV immunization reduced PRRSV-2 viraemia but did not reduce PRRSV-2 replication in the respiratory tract after challenge. PRRSV-2/H3N2 co-infection did not alter the effect of PRRS MLV vaccination on PRRSV-2 load. However, in unimmunized animals, co-infection had a beneficial effect by reducing PRRSV-2 viraemia and H3N2 shedding, with a trend for lower PRRSV-2 and H3N2 loads in the BALF. However, the reduction of H3N2 viral load was reversed in PRRS MLV immunized animals.

### PRRSV-2-specific antibody responses

We next determined whether PRRSV-2/H3N2 co-infection modulates the PRRS MLV induced Ab responses. Ab in serum were evaluated using a commercial PRRSV N-specific ELISA at 0 dpv, 20 dpv and 5 dpc (38 dpv) (**Figure 3A**). Sera from immunized pigs at 20 dpv were above the cut-off and displayed significant levels of PRRSV N-specific Abs (p<0.05; 0 vs 20 dpv) indicating a seroconversion following PRRS MLV immunization. There was no significant difference in the level of PRRSV N-specific Abs between the Vac groups at 20 dpv. However, after the challenge, an increase of PRRSV N-specific Abs was detected in the Vac + PRRSV-2/H3N2 group (p<0.05; 20 vs 38 dpv) but not in the Vac + PRRSV-2 group. The levels of PRRSV N-specific antibodies were comparable between Vac + PRRSV-2 and Vac + PRRSV-2/H3N2 groups at 5 dpc (p=0.13).

**Figure 3:**
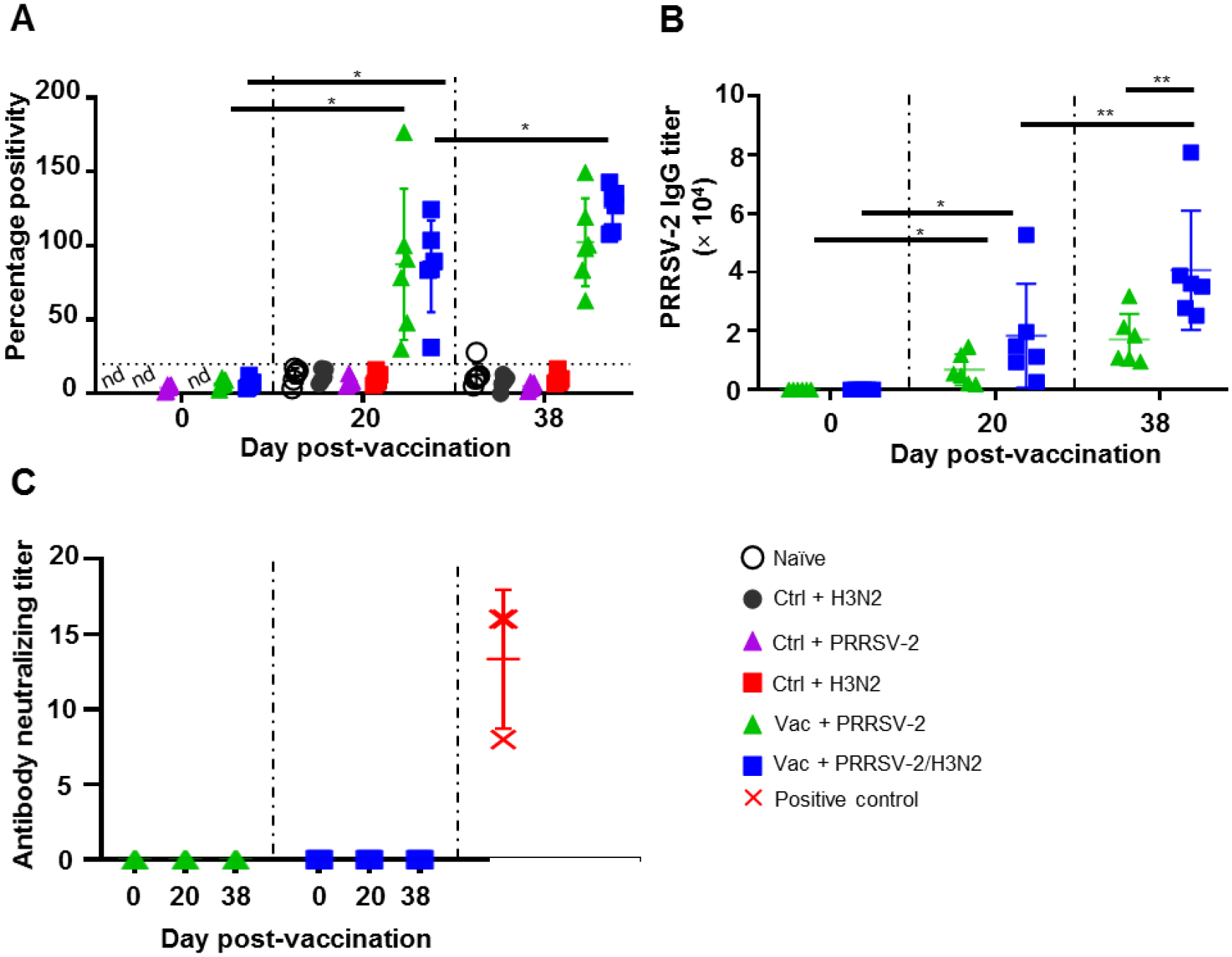
Antibody responses against PRRSV-2. (**A**) Detection of PRRSV N-specific Abs in the serum at 0, 20 and 38 dpv was performed with a commercial ELISA test. The positive threshold is indicated with dashed lines. (**B**) PRRSV-specific Ab titers were measured in the sera of PRRS-immunized pigs at 0, 20 and 38 dpv with an-in house ELISA test. (**C**) Levels of virus neutralizing Ab titer in the serum of PRRS-vaccinated pig at 0, 20 and 38 are shown. Sera from PRRSV-2 infected pigs from an unrelated study were used as positive controls (red symbols). Each pig serum is shown as a symbol within the indicated group (n=6 per group) and the mean ± SD is represented. The comparison between the percentage positivity values or Ab titers measured at 20 and 38 dpv versus at 0 dpv for each vaccinated group were performed using the Wilcoxon test. Comparisons between groups were made using the Mann-Whitney test. Asterisks indicate significant differences (*p < 0.05; **p < 0.01). nd: not determined.

Serum Ab titers were also measured by an in-house ELISA using lysate from PRRSV-2 16CB02 infected cells as antigens (**Figure 3B**). Consistent with the results found with the N-based ELISA test, significant anti-PRRSV-2 IgG titers were measured after immunization (p<0.05; 0 vs 20 dpv) which were similar in both Vac + PRRSV-2 and Vac + PRRSV-2/H3N2 groups (p=0.13; 20 dpv). However, after challenge, the Vac + PRRSV-2/H3N2 group had significantly higher Ab titers compared to the Vac + PRRSV-2 group (mean Ab titer: 40,647 vs 17,126, respectively; p<0.01), suggesting that the co-infection with H3N2 enhanced the recall of B cell responses primed by PRRS MLV immunization. Remarkably, none of the sera collected at 20 and 38 dpv were able to neutralize the PRRSV-2 16CB02 challenge strain (**Figure 3C**). The lack of neutralization might be explained by differences between the vaccine and challenge strains (**Supplementary Table S4**).

These data indicated that PRRS MLV vaccination induced a significant PRRSV-2-specific Ab response, which was enhanced after PRRSV-2/H3N2 co-infection. In contrast, the PRRSV-2 only challenge did not significantly alter the magnitude of the specific antibody response (**Figure 3C**).

### PRRSV-2-specific T cell responses in PBMC

In swine, conventional CD4^+^ T cells are defined as CD3^+^CD4^+^ CD8α^+/-^CD8β^-^ and CD8^+^ T cells as CD3^+^CD4^-^CD8β^+^ (36, 37). PRRSV-2 specific CD4^+^ and CD8^+^ T cell responses were assessed by intracellular staining of IFN-γ, TNF, IL-2, IL-4, and IL-17 after *in vitro* restimulation of PBMC isolated at 0, 20 and 38 dpv (5 dpc) with PRRSV-2 (**Supplementary Figures S3 and S4**). Overall, the frequencies of cytokine-producing cells were low in T cell populations after PRRS MLV immunization, as demonstrated previously (11) (**Figure 4**). Similar results were obtained after the challenge with either PRRSV-2 alone or together with H3N2, although a substantial increase in IFN-γ expressing CD8^+^ T cells was detected after in the Vac + PRRSV-2/H3N2 group but not the Vac + PRRSV-2 group (mean 0.11 % and 0.06 %, respectively) (**Figure 4B**). Similarly, an increased proportion of PRRSV-2-specific TNF expressing CD8^+^ T cell population was detected in these two groups (mean 0.24 % and 0.35 %), but this was not boosted after challenge. In addition, no IL-2, IL-4 and IL-17 expressing CD4^+^ and IL-2 expressing CD8^+^ T cells were detected after challenge (**Figure 4** and **supplementary Figure S5**).

**Figure 4:**
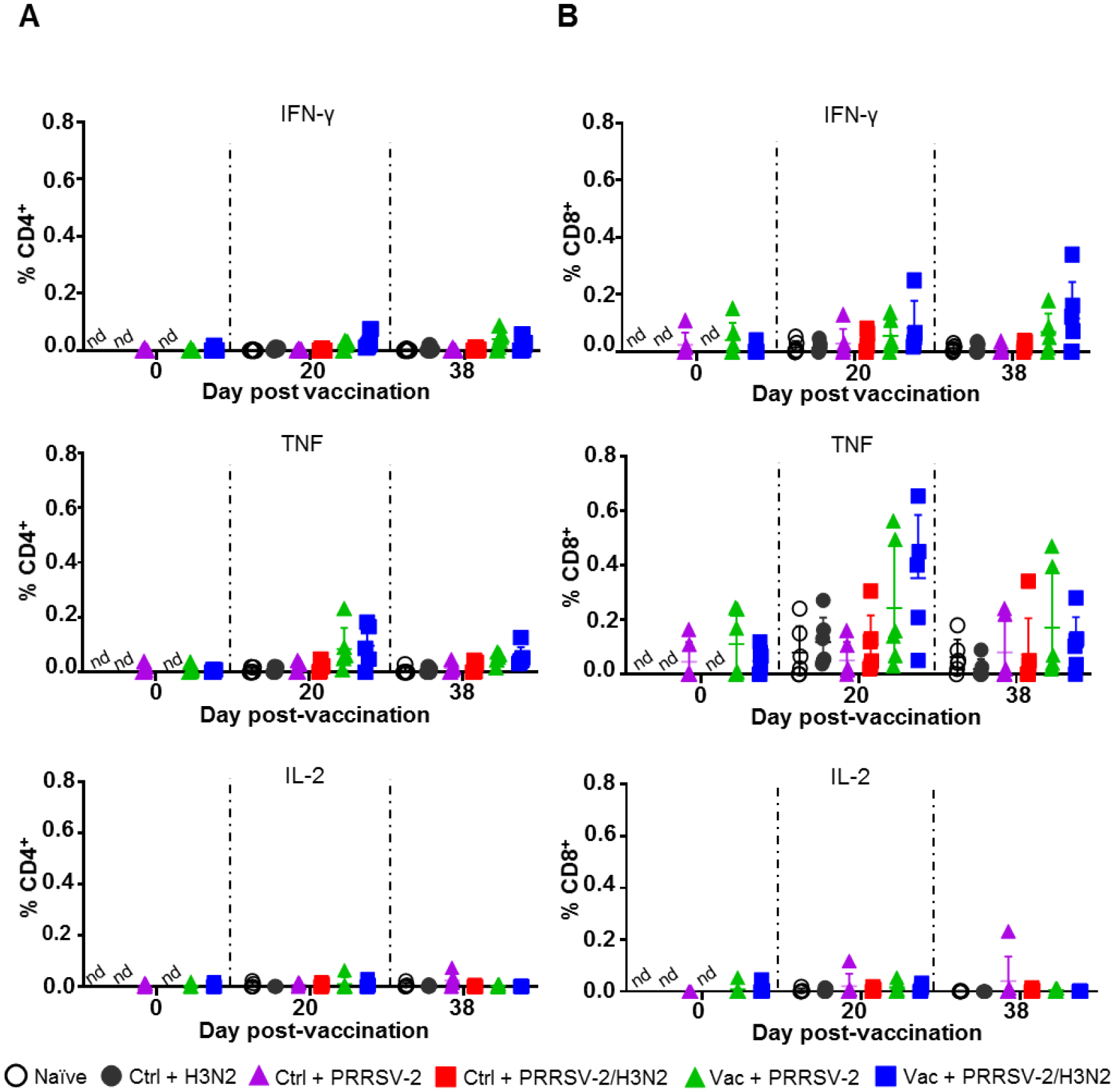
PRRSV-2-specific T cell responses in PBMC. PBMC isolated at 0, 20 and 38 dpv were restimulated *in vitro* for 18h with PRRSV-2 (MOI 0.1) or cultured with medium. Intracellular staining of IFN-γ, TNF, IL-2, was performed and frequencies of IFN-γ, TNF and IL-2 producing CD4^+^ (**A**) and CD8β^+^ (**B**) T cells were analyzed. The corrected frequencies values are shown (percentage of cytokine-producing cells subtracted with medium only). Data for individual pigs and the group mean ± SD are displayed (n=5-6 per group). The Wilcoxon test was used to compare the T cell responses at day 0 and 20 dpv within the same group. Comparisons between 2 groups were performed using Mann-Whitney test. nd: not determined.

Overall, PRRS MLV immunization induced TNF and IFN-γ expressing CD8^+^ T cells in PBMC, which were not boosted by the subsequent PRRSV-2 or PRRSV-2/H3N2 challenge. PRRSV-2-specific CD4^+^ T cell responses were weaker compared to CD8^+^ T cells and there was no significance between the groups for any of the measured cytokines.

### PRRSV-2- and H3N2-specific T cell responses in BALF

Local T cell responses in the lung likely to play a role in controlling both IAV (38–40) and PRRSV-2 infections and pathology (11, 41). We therefore assessed the αβ and γδ T cell responses against both PRRSV-2 and H3N2 in BALF at 5 dpc. Cells were restimulated with PRRSV-2 or H3N2, and cytokine production was assessed by ICS. Staining of IFN-γ, TNF, IL-2 were performed for CD4^+^ and CD8^+^ αβ T cells, and staining of IFN-γ, TNF, IL-17 for γδ T cells (**Supplementary Figure S6**), which were divided into CD2^+^ and CD2^-^ subsets (42). CD4^+^ and CD8^+^ T cell responses to PRRSV-2 overall were very low after single infection with PRRSV-2 in either Ctrl or Vac groups (**Figure 5A**). However, T cell responses to PRRSV-2 stimulation were higher after PRRSV-2/H3N2 co-infection in both Ctrl and Vac pigs. The highest proportion of IL-2^+^ CD4^+^ (mean 0.14 %) and IFN-γ^+^ CD8^+^ T cells (mean 0.11 %) was observed in Ctrl + PRRSV-2/H3N2 group. In the Vac group, PRRSV-2/H3N2 co-infection induced a proportion of IFN-γ^+^ CD8^+^ T cells (mean 0.23 %) and a significantly greater proportion of TNF^+^ CD8^+^ T cells compared to Vac + PRRSV-2 (mean 0.35 %; p<0.5). Similarly, co-infection induced the highest proportion of IFN-γ^+^ CD2^+^ γδ T cells in the Vac group (mean 0.22 %), and IL-17^+^ CD2^+^ γδ T cells in the Ctrl group (mean 0.30 %) (**Figure 5B**). Higher frequencies of TNF^+^ and IL-17^+^ CD2^-^ γδ T cells were detected in the Ctrl + PRRSV-2/H3N2 compared to Ctrl + PRRSV-2 group (mean 0.73 % versus 0.31 % and mean 0.40 % versus 0.23 %, respectively) though this was not statistically significant. In vaccinated pigs, co-infection increased the frequency of TNF^+^ (mean 0.47 %) and IL-17^+^ (mean 0.51 %) CD2^-^ γδ T cells, although these differences did not reach statistical significance.

**Figure 5:**
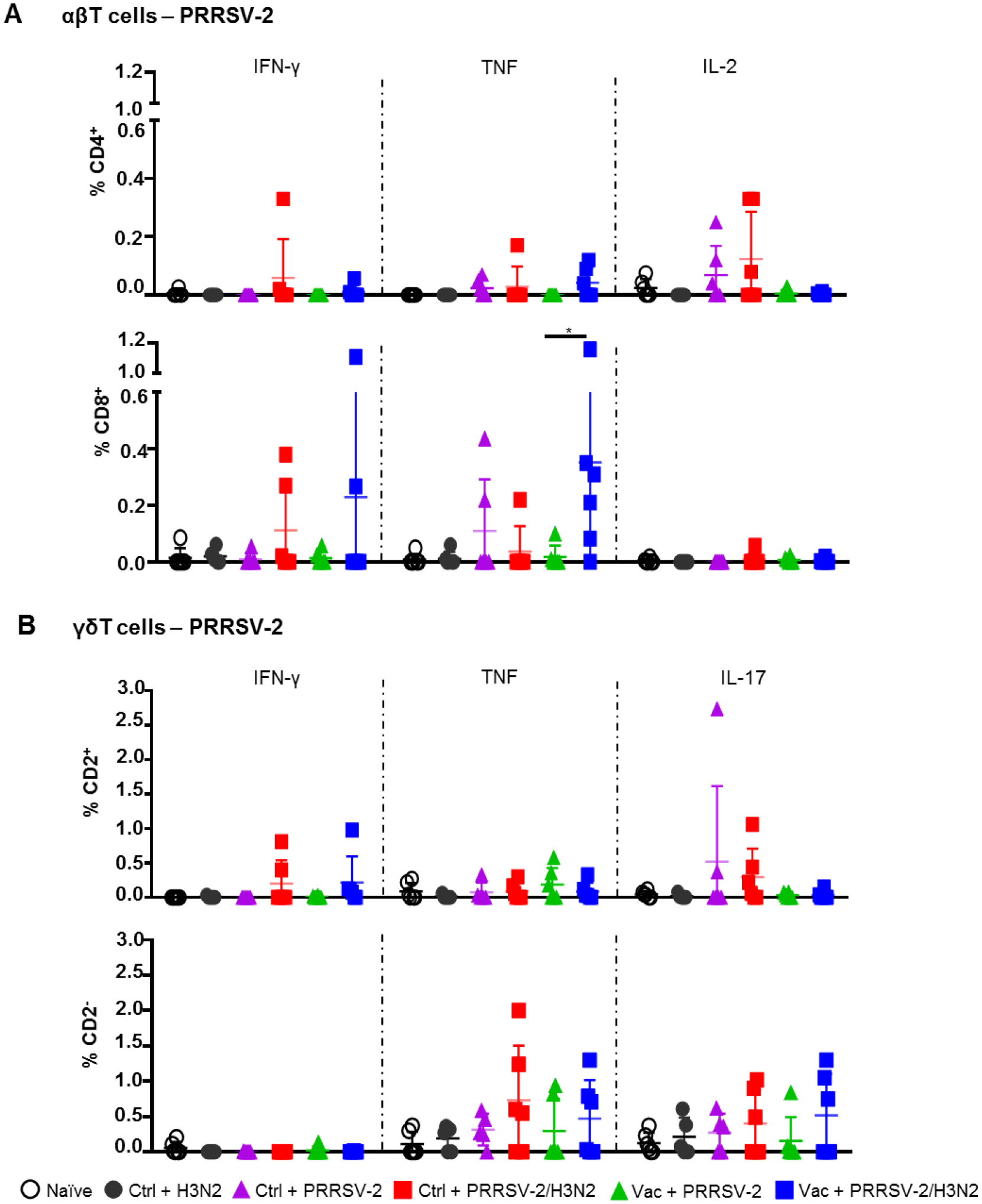
PRRSV-2-specific T cell responses in bronchoalveolar lavage. Cells isolated from BAL at 5 dpc were restimulated with PRRSV-2 (MOI 0.1) or cultured with medium. (**A**) Frequency of IFN-γ-, TNF- and IL-2-producing CD4^+^ and CD8β^+^ T cells are shown. (**B**) Frequency of IFN-γ-, TNF- and IL-17-producing CD2^+^ and CD2^-^ γδ T cells are represented. The corrected frequencies (percentage of cytokine-producing cells subtracted with medium only) of each individual pig and the group mean ± SD are displayed (n=5-6 per group). Comparisons between groups were made using Mann-Whitney test. Asterisk indicates significant difference (*p < 0.05).

Stimulation of BALC with H3N2 also showed the greatest frequencies of cytokine producing T cells in co-infected animals (**Figure 6A**). The proportion of IL-2 expressing CD4^+^ T cells in the Ctrl + PRRSV-2/H3N2 group was higher compared to Ctrl + H3N2 (mean 0.23 % versus 0.05 %). Similarly, the frequency of IFN-γ^+^ CD8^+^ T cells in the Ctrl + PRRSV-2/H3N2 group was higher compared to the Ctrl + H3N2 (mean 0.35 % versus 0.07 %), but none of these differences reached statistical significance. A greater frequency of TNF**^+^** CD2^+^ γδ T cells was again measured in the Ctrl + PRRSV-2/H3N2 group compared to Ctrl + H3N2 group (mean 0.29 % versus 0.01 %) (**Figure 6B**). Similar results were obtained for TNF^+^ and IL-17^+^ CD2^-^ γδ T cells in the Ctrl + PRRSV-2/H3N2 group compared to the Ctrl + H3N2 group (mean 1.18 % versus 0.23 % and 1.02 % versus 0.28 %, respectively). Intriguingly, lower H3N2-specific cytokine producing T cell responses were induced by the co-infection in PRRS MLV vaccinated group in comparison to the unvaccinated group although no statistically significant differences were observed (**Figures 6A and 6B**).

**Figure 6:**
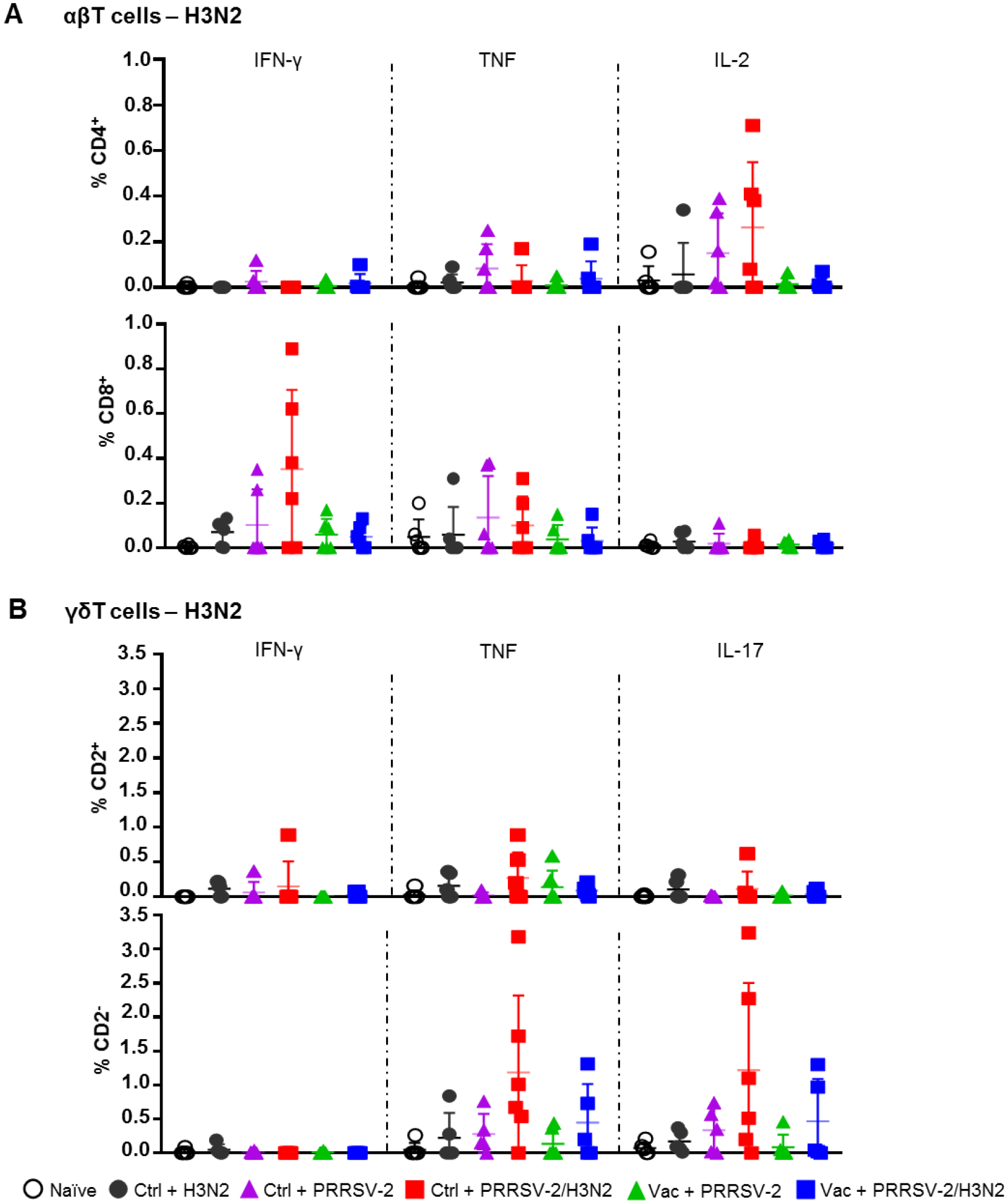
H3N2-specific T cell responses in bronchoalveolar lavage. Cells isolated from BALF at 5 dpc were restimulated with swIAV H3N2 (MOI 0.1) or cultured with medium. Cytokine secretion measured in CD4^+^ and CD8β^+^ (**A**), and CD2^+^ and CD2^-^ γδ (**B**) T cells are represented. Corrected frequencies of individual value and the mean ± SD are displayed (n=5-6 per group). Comparisons between 2 groups were made using the Mann-Whitney test.

Overall, these data indicate that PRRSV-2/H3N2 co-infection induces the highest frequency of cytokine producing T cells in response to both PRRSV-2 and H3N2 in BAL, although these did not reach statistical significance, most likely due to the sample size. The data also highlight the effect of PRRS MLV vaccination on the immune responses after the co-infection. The increased PRRSV-specific, but lowered H3N2-specific T cell responses in the Vac + PRRSV-2/H3N2 compared to the Ctrl + PRRSV-2/H3N2 suggest that PRRS MLV may potentially drive the host immune response toward PRRSV-specific, and rather away from the H3N2-specific responses.

### Cytokine expression in the lung

To further characterize the immune responses in the lungs of singly and co-infected pigs, the gene expression of a panel of cytokines and chemokines was assessed in lung tissues and data were normalized to the naïve group (**Figure 7**). After single infection with H3N2, an elevated mRNA expression of pro-inflammatory cytokines TNF and IFN-γ (mean fold change of 2.22 and 1.86, respectively), and anti-inflammatory cytokines IL-10 and TGF-β (mean fold change of 1.92 and 2.40 respectively), was observed (**Figure 7**). In PRRSV-2-infected group (Ctrl + PRRSV-2), a modest increase of TNF, IL-12p40 and IL-10 transcripts (mean fold change of 1.67, 1.69 and 1.80, respectively) was measured. After co-infection (Ctrl + PRRSV-2/H3N2), TNF, IFN-γ, IL-12p40, IL-4 and CXCL-13 transcripts were all up-regulated (mean fold change of 2.43, 1.86, 3.47, 2.49 and 2.83, respectively). In both vaccinated single and co-infected groups, an up-regulation of TNF (mean fold change 2.0 in both Vac groups) and IFN-γ transcripts were quantified (mean fold change 1.8 in both Vac groups). Notably both IL-10 (mean fold change of 2.5 and 2.4, respectively) and TGF-β (mean fold change of 1.6 and 2.0, respectively) mRNA expression was also upregulated in these vaccinated groups. A significant down-regulation of IL-12p40 mRNA was observed in Vac + PRRSV-2/H3N2 pigs compared to the Ctrl + PRRSV-2/H3N2 group (p<0.05).

**Figure 7:**
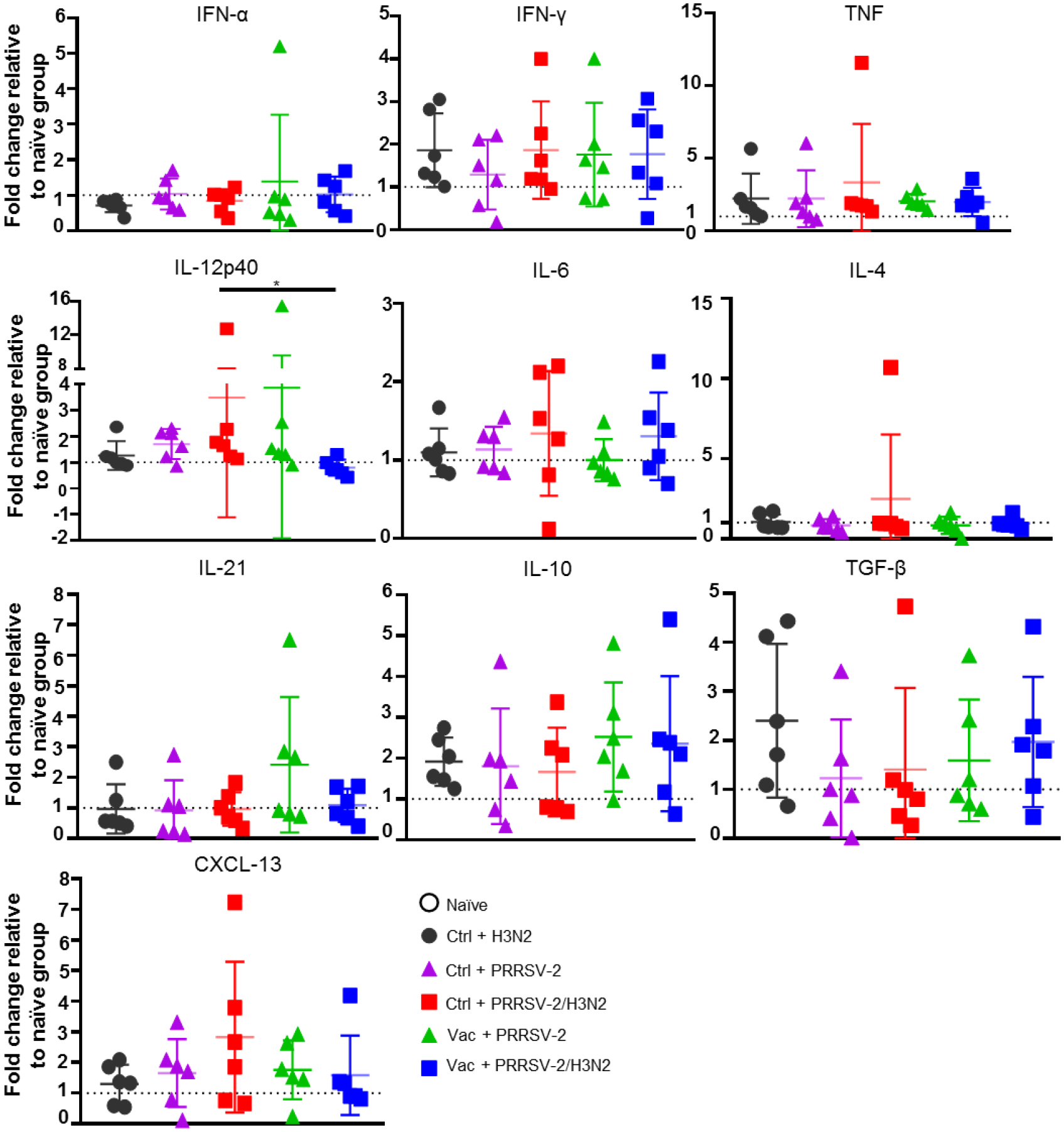
Gene expression in lung tissues. Total RNA was extracted from lung tissue collected at 5 dpc and the relative mRNA expression of IFN-α, IFN-γ, TNF, IL-12p40, IL-4, IL-6, IL-21, IL-10, TGF-β and CXCL-13 was assessed by qRT-PCR. Fold changes are shown over naïve group (dash line) after normalization with *GAPDH* and *RPS24* genes. Individual pig values and the group mean ± SD are displayed (n=5-6 per group). Comparisons were made using Kruskal-Wallis test and asterisks indicate significant differences (*p < 0.05).

Despite a lack of statistical significance, these data suggest that upregulation of inhibitory cytokines gene expression in the lungs of previously immunized pigs might have influenced responses to H3N2 and PRRSV-2.

## Discussion

The PRDC is responsible for major economic losses in the pig industry worldwide. The PRDC commonly results from mixed infections, in combination with environmental stressors (16). PRRSV and swIAV, alone or in combination, are two major viral pathogens involved in the PRDC, often leading to secondary infections with opportunistic bacteria. Previous experimental *in vivo* PRRSV/swIAV co-infection studies reported variable outcomes in terms of clinical or virological parameters, which may reflect differences in the timing of infections (18,43,44). Superinfection, i.e., infection of pigs with H1N1 3 days later a primary infection with PRRSV-1 led to more severe disease and a delayed shedding of H1N1 compared to pigs that were infected 14 days later (45), whereas simultaneous PRRSV-1/H1N1 co-infection did not alter the clinical and virological course of infection of either virus (44). Several studies have also assessed the effect of concurrent infections and superinfections on vaccines efficacy and demonstrated that PRRSV infection decreases the efficacy of swIAV (46), *Mycoplasma hyopneumoniae* (47), and classical swine fever (48) vaccines although underlying mechanisms involved have not been demonstrated. Whilst vaccination with PRRS MLV is widely practiced in efforts to control PRRS, swIAV vaccination is less often used. Reflecting the field situation, a recent study experimentally assessed the impact of swIAV superinfection on vaccination with PRRS MLV efficacy and found a delay in MLV replication and Ab responses, but this did not affect vaccine efficacy (19). In our study here, we investigated the effect of PRRSV-2/H3N2 co-infection on the protection provided by a commercial PRRS MLV vaccine. The PRRS MLV protected against clinical disease, reduced lung pathology and viremia, but not virus shedding, as previously reported (14,49–51). However, PRRSV-2/H3N2 co-infection abrogated the protective effect of PRRS MLV vaccination on clinical disease and pathology. Co-infection did not affect the vaccine-induced reduction in PRRSV-2 load and enhanced CD8^+^ T cell responses in the lung and Ab responses. In contrast, co-infection in non-immunized animals had a beneficial effect by reducing PRRSV-2 viremia and H3N2 virus loads in BALF and nasal swabs at day 5 post-infection and did not affect clinical signs or pathology.

We sought to dissect the mechanisms underlying these opposing effects of co-infection in immunized and non-immunized animals at day 5 post-infection. The reduced PRRSV load in unimmunized co-infected animals has been suggested to be due to the early type I IFN response triggered by H3N2, and interference and/or competition for substrates required for replication (19, 52). In this study, analysis of gene expression in the lungs, did not reveal significant differences in the expression of IFN-α, which may be due to the timing of sampling (5 days post-infection). However, there was significantly more IL-12p40 gene expression, in the lung, and a trend for higher number of H3N2-specific T cell responses (IL-2^+^ CD4^+^, IFN-γ^+^ CD8^+^, IL-17^+^ CD2^-^ γδ and TNF^+^ CD2^-^ γδ T cells) and PRRSV-2-specific T cell responses (IL-17^+^ CD2^-^ γδ T cells TNF^+^ CD2^-^ γδ T cells) in the BALF of the co-infected unimmunized animals. This trend for enhanced cell-mediated response, may have contributed to the control of virus replication in the lung in the co-infected groups compared to the singly infected groups. In vaccinated pigs, co-infection induced a greater PRRSV-specific TNF^+^ CD8^+^ T cell response in the BALF and an increased Ab response. However, this was not the case for the specific response to H3N2, and the cytokine gene expression profile in the lung did not differ significantly from the singly infected animals. The precise mechanism for abrogation of the PRRS MLV protective effect against lung injury by co-infection remains unclear but might be due to exuberant cytokine production by other cell types producing more pro-inflammatory cytokines such as TNF and IL-6 (53), which were not detected at the timepoint sampled here. Although effectors CD8^+^ T cells in the BAL aids to eliminate PRRSV-infected cells, the local response might contribute to pulmonary inflammation and injury. Alternatively, co-infection may have enhanced the production of low affinity non-neutralizing antibodies that may augment infection and exacerbate disease.

Whilst PRRS MLV vaccines can provide a significant clinical benefit, the protection against virus shedding is limited which may drive the evolution of PRRSV (54, 55). This has been attributed to these vaccines being weakly immunogenic, especially for cellular responses (8). In line with previous reports, we confirmed that PRRS MLV vaccinated pigs recorded lower lung lesions, viraemia and viral load in lungs after a single infection with PRRSV-2, which was associated with a robust, although non-neutralizing, antibody response and a weak peripheral PRRSV-specific T cell response (11,14,50,51).

PRRSV has been shown to suppress host immune response through the induction of regulatory T cells (56), alteration of peripheral NK cell cytotoxic activity (57), and inhibition of cytokine responses (58, 59). Moreover, PRRS MLV vaccine can induce systemic secretion of IL-10 (60). It is therefore plausible that the vaccine used in our study may exhibit similar immunomodulatory features. We observed that, in comparison to unvaccinated pigs, lungs from vaccinated pigs co-infected with PRRSV-2/H3N2 displayed a lower level of pro-inflammatory IL-12p40 gene expression, along with a trend for higher expression of anti-inflammatory IL-10 and TGF-β as measured by RT-qPCR. In a study on *Litomosoides sigmodontis* helminth infection (60), helminth-infected mice exhibit lower HA-specific antibody responses post-IAV vaccination, linked with a higher viral load in the lung compared to non-infected mice. The level of HA-specific induced by the vaccine was restored after the blockade of IL-10 using an anti-IL-10 receptor mAb. PRRS MLV vaccination may have potentially impaired the host immune response to H3N2 and perhaps facilitated its replication. In addition to the highest lung pathology seen in the PRRS MLV vaccinated co-infected pigs, the beneficial effect of the co-infection in reducing H3N2 viral loads was also abrogated in vaccinated pigs. These data suggests that PRRS MLV initiates a strong PRRS specific response and may suppress H3N2 specific responses leading to increased pathology and poor disease outcome.

Whilst confirmatory studies are required, the demonstration that co-infection with H3N2 can abrogate the clinical protective effect of PRRSV MLV suggests that better control measures against swIAV might be warranted to further increase protection against PRRSV. Large number of herds are endemically infected with swIAV and suffer intermittent bouts of disease. SwIAV contributes to suboptimal weight gain and reproductive performance and is occasionally associated with fever-induced abortion in sows (61). Immunization can be a cost-effective control measure to combat swIAV, but the rapid evolution of the virus is a major obstacle (62). Not all swIAV-endemic countries use vaccines to control the disease, for example, current UK policy does not involve immunization against swIAV, although it is used in some European countries and widely in the USA (63). We have used recent field PRRS and swIAV strains from Thailand. As in the UK, immunization of pigs against swIAV is not mandatory in Thailand, although some farms use the GRIPORK^®^ vaccine (Hipra, Spain) containing inactivated H1N1 and H3N2 strains. Inactivated vaccines induce neutralizing Ab against the immunizing strain but do not induce sufficient heterologous protection due to the rapid viral escape that occurs through antigenic drift of the surface glycoproteins (64), so that coincident infection with PRRS and swIAV remains possible. There is a need to develop more broadly protective influenza vaccines that could provide a better control of swIAV, reducing the zoonotic risk (65), the contribution to the PRDC and its potentially harmful effect on PRRS vaccine efficacy.

## Data availability

The raw data supporting the conclusions of this article will be made available by the authors, without undue reservation.

## Competing interests

*The authors declare that the research was conducted in the absence of any commercial or financial relationships that could be construed as a potential conflict of interest*.

## Funding

This study was supported by the Newton Fund UK-China-Philippines-Thailand Swine and Poultry Research Initiative award BB/R01275X/1 (Broadly protective vaccines for porcine reproductive and respiratory syndrome and swine influenza virus infections), UKRI Biotechnology and Biological Sciences Research Council (BBSRC) Institute Strategic Programme and Core Capability Grants to The Pirbright Institute (BBS/E/I/00007031 and BBS/E/I/00007037 and BBS/E/I/00007039).

## Acknowledgements

The authors would like to acknowledge The Pirbright Institute Flow Cytometry Unit, and Animal Services Team for animal care and their invaluable help for providing samples. We would also like to thank Miriam Pedrera and Rebecca K. McLean for their assistance during post-mortems.

## Author Contributions

ET, SG, NW and SK acquired funding for the project. ET, SG and TC contributed to the conception, design, and coordination of the study. TC and EM performed experiments. TC acquired, analyzed, and interpreted the data. ET, SG, EM, EV, VM, BP, AM, TM, and ME contributed to sampling during post-mortem. FS carried out pathological analysis. TC wrote the first draft of the manuscript. ET, SG, NW, and SK edited and revised the manuscript. All authors approved the submitted version.

## Notes

### Competing Interest Statement

The authors have declared no competing interest.

## References

1. Holtkamp DJ, Kliebenstein JB, Neumann EJ, Zimmerman JJ, Rotto HF, Yoder TK, et al. Assessment of the economic impact of porcine reproductive and respiratory syndrome virus on United States pork producers. J Swine Heal Prod. (2013) 21:72–84.

2. Stadejek T, Larsen LE, Podgórska K, Bøtner A, Botti S, Dolka I, et al. Pathogenicity of three genetically diverse strains of PRRSV Type 1 in specific pathogen free pigs. Vet Microbiol. (2017) 209:13–19. doi:10.1016/j.vetmic.2017.05.011

3. Weesendorp E, Rebel JMJ, Popma-De Graaf DJ, Fijten HP, Stockhofe-Zurwieden N. Lung pathogenicity of European genotype 3 strain porcine reproductive and respiratory syndrome virus (PRRSV) differs from that of subtype 1 strains. Vet Microbiol. (2014) 174:127–38. doi:10.1016/j.vetmic.2014.09.010

4. Montaner-Tarbes S, del Portillo HA, Montoya M, Fraile L. Key Gaps in the Knowledge of the Porcine Respiratory Reproductive Syndrome Virus (PRRSV). Front Vet Sci. (2019) 6:38. doi:10.3389/fvets.2019.00038

5. Gebhardt JT, Tokach MD, Dritz SS, DeRouchey JM, Woodworth JC, Goodband RD, et al. Postweaning mortality in commercial swine production. I: Review of non-infectious contributing factors. Transl Anim Sci. (2020) 4: 485–506. doi:10.1093/tas/txaa052

6. Pejsak Z, Stadejek T, Markowska-Daniel I. Clinical signs and economic losses caused by porcine reproductive and respiratory syndrome virus in a large breeding farm. Vet Microbiol. (1997) 55:317–22. doi:10.1016/s0378-1135(96)01326-0

7. Walker PJ, Siddell SG, Lefkowitz EJ, Mushegian AR, Adriaenssens EM, Alfenas-Zerbini P, et al. Changes to virus taxonomy and to the International Code of Virus Classification and Nomenclature ratified by the International Committee on Taxonomy of Viruses (2021). Arch Virol. 2021 (2021) 166: 2633–48. doi:10.1007/s00705-021-05156-1

8. Zhou L, Ge X, Yang H. Porcine reproductive and respiratory syndrome modified live virus vaccine: A “leaky” vaccine with debatable efficacy and safety. Vaccines. (2021) 9:362. doi:10.3390/vaccines9040362

9. Nan Y, Wu C, Gu G, Sun W, Zhang Y-J, Zhou E-M. Improved Vaccine against PRRSV: Current Progress and Future Perspective. Front Microbiol. (2017) 8:1635. doi:10.3389/fmicb.2017.01635

10. Geldhof MF, Vanhee M, Van Breedam W, Van Doorsselaere J, Karniychuk UU, et al. Comparison of the efficacy of autogenous inactivated Porcine Reproductive and Respiratory Syndrome Virus (PRRSV) vaccines with that of commercial vaccines against homologous and heterologous challenges. BMC Vet Res. (2012) 8:182. doi:10.1186/1746-6148-8-182

11. Kick AR, Amaral AF, Cortes LM, Fogle JE, Crisci E, Almond GW, et al. The T-Cell Response to Type 2 Porcine Reproductive and Respiratory Syndrome Virus (PRRSV). Viruses. (2019) 11:796. doi:10.3390/v11090796

12. Osorio FA, Galeota JA, Nelson E, Brodersen B, Doster A, Wills R, et al. Passive transfer of virus-specific antibodies confers protection against reproductive failure induced by a virulent strain of porcine reproductive and respiratory syndrome virus and establishes sterilizing immunity. Virology. (2002) 302:9–20. doi:10.1006/viro.2002.1612

13. Lopez OJ, Oliveira MF, Alvarez Garcia E, Kwon BJ, Doster A, Osorio FA. Protection against Porcine Reproductive and Respiratory Syndrome Virus (PRRSV) infection through passive transfer of PRRSV-neutralizing antibodies is dose dependent. Clin Vaccine Immunol. (2007) 14:269–75. doi:10.1128/CVI.00304-06

14. Kick AR, Amaral AF, Frias-De-Diego A, Cortes LM, Fogle JE, Crisci E, et al. The Local and Systemic Humoral Immune Response Against Homologous and Heterologous Strains of the Type 2 Porcine Reproductive and Respiratory Syndrome Virus. Front Immunol. (2021) 12:637613. doi:10.3389/fimmu.2021.637613

15. Rahe MC, Murtaugh MP. Mechanisms of adaptive immunity to porcine reproductive and respiratory syndrome virus. Viruses. (2017) 9:148. doi:10.3390/v9060148

16. Saade G, Deblanc C, Bougon J, Marois-Créhan C, Fablet C, Auray G, et al. Coinfections and their molecular consequences in the porcine respiratory tract. Vet Res. (2020) 51:80. doi:10.1186/s13567-020-00807-8

17. Choi YK, Goyal SM, Joo HS. Retrospective analysis of etiologic agents associated with respiratory diseases in pigs. Can Vet J. (2003) 44:735–37.

18. Van Reeth K, Nauwynck H, Pensaert M. Dual infections of feeder pigs with porcine reproductive and respiratory syndrome virus followed by porcine respiratory coronavirus or swine influenza virus: A clinical and virological study. Vet Microbiol. (1996) 48:325–35. doi:10.1016/0378-1135(95)00145-x

19. Renson P, Deblanc C, Bougon J, Le Dimna M, Gorin S, Mahé S, et al. Concomitant swine influenza a virus infection alters PRRSV1 MLV viremia in piglets but does not interfere with vaccine protection in experimental conditions. Vaccines. (2021) 9:356. doi:10.3390/vaccines9040356

20. Hoffmann E, Stech J, Guan Y, Webster RG, Perez DR. Universal primer set for the full-length amplification of all influenza A viruses. Arch Virol. (2001) 146:2275–89. doi:10.1007/s007050170002

21. Ryt-Hansen P, Pedersen AG, Larsen I, Krog JS, Kristensen CS, Larsen LE. Acute Influenza A virus outbreak in an enzootic infected sow herd: Impact on viral dynamics, genetic and antigenic variability and effect of maternally derived antibodies and vaccination. PLoS One. (2019) 14:e0224854. doi:10.1371/journal.pone.0224854

22. Holzer B, Morgan SB, Martini V, Sharma R, Clark B, Chiu C, et al. Immunogenicity and Protective Efficacy of Seasonal Human Live Attenuated Cold-Adapted Influenza Virus Vaccine in Pigs. Front Immunol. (2019) 10:2625. doi:10.3389/fimmu.2019.02625

23. Morgan SB, Frossard JP, Pallares FJ, Gough J, Stadejek T, Graham SP, et al. Pathology and Virus Distribution in the Lung and Lymphoid Tissues of Pigs Experimentally Inoculated with Three Distinct Type 1 PRRS Virus Isolates of Varying Pathogenicity. Transbound Emerg Dis. (2016) 63:285–95. doi:10.1111/tbed.12272

24. Morgan SB, Hemmink JD, Porter E, Harley R, Shelton H, Aramouni M, et al. Aerosol Delivery of a Candidate Universal Influenza Vaccine Reduces Viral Load in Pigs Challenged with Pandemic H1N1 Virus. J Immunol. (2016) 196:5014–23. doi:10.4049/jimmunol.1502632

25. Gauger PC, Vincent AL, Loving CL, Henningson JN, Lager KM, Janke BH, et al. Kinetics of Lung Lesion Development and Pro-Inflammatory Cytokine Response in Pigs With Vaccine-Associated Enhanced Respiratory Disease Induced by Challenge With Pandemic (2009) A/H1N1 Influenza Virus. Vet Pathol. (2012) 49:900–12. doi:10.1177/0300985812439724

26. McNee A, Smith T, Holzer B, Clark B, Bessell E, Guibinga G, et al. Establishment of a Pig Influenza Challenge Model for Evaluation of Monoclonal Antibody Delivery Platforms. J Immunol. (2020) 205:648–60. doi:10.4049/jimmunol.2000429

27. Bordet E, Blanc F, Tiret M, Crisci E, Bouguyon E, Renson P, et al. Porcine reproductive and respiratory syndrome virus type 1.3 Lena triggers conventional dendritic cells 1 activation and t helper 1 immune response without infecting dendritic cells. Front Immunol. (2018) 9:2299. doi:10.3389/fimmu.2018.02299

28. Liu G, Wang Y, Jiang S, Sui M, Wang C, Kang L, et al. Suppression of lymphocyte apoptosis in spleen by CXCL13 after porcine circovirus type 2 infection and regulatory mechanism of CXCL13 expression in pigs. Vet Res. (2019) 50:17. doi:10.1186/s13567-019-0634-2

29. Vandesompele J, De Preter K, Pattyn F, Poppe B, Van Roy N, De Paepe A, et al. Accurate normalization of real-time quantitative RT-PCR data by geometric averaging of multiple internal control genes. Genome Biol. (2002) 3:research0034. doi:10.1186/gb-2002-3-7-research0034

30. Maisonnasse P, Bouguyon E, Piton G, Ezquerra A, Urien C, Deloizy C, et al. The respiratory DC/macrophage network at steady-state and upon influenza infection in the swine biomedical model. Mucosal Immunol. (2016) 9:835–49. doi:10.1038/mi.2015.105

31. Chrun T, Lacôte S, Urien C, Richard CA, Tenbusch M, Aubrey N, et al. A DNA vaccine encoding the Gn ectodomain of Rift valley fever virus protects mice via a humoral response decreased by DEC205 targeting. Front Immunol. (2019) 10:860. doi:10.3389/fimmu.2019.00860

32. Robinson SR, Li J, Nelson EA, Murtaugh MP. Broadly neutralizing antibodies against the rapidly evolving porcine reproductive and respiratory syndrome virus. Virus Res. (2015) 203:56–65. doi:10.1016/j.virusres.2015.03.016

33. Martini V, Paudyal B, Chrun T, McNee A, Edmans M, Atangana Maze E, et al. Simultaneous Aerosol and Intramuscular Immunization with Influenza Vaccine Induces Powerful Protective Local T Cell and Systemic Antibody Immune Responses in Pigs. J Immunol. (2021) 206:652–63. 10.4049/jimmunol.2001086

34. Lyoo KS, Kim JK, Jung K, Kang BK, Song D. Comparative pathology of pigs infected with Korean H1N1, H1N2, or H3N2 swine influenza A viruses. Virol J. (2014) 11:170. doi:10.1186/1743-422X-11-170

35. Halbur PG, Paul PS, Frey ML, Landgraf J, Eernisse K, Meng XJ, et al. Comparison of the pathogenicity of two US porcine reproductive and respiratory syndrome virus isolates with that of the Lelystad virus. Vet Pathol. (1995) 32:648–60. doi:10.1177/030098589503200606

36. Edmans M, McNee A, Porter E, Vatzia E, Paudyal B, Martini V, et al. Magnitude and Kinetics of T Cell and Antibody Responses During H1N1pdm09 Infection in Inbred Babraham Pigs and Outbred Pigs. Front Immunol. (2021) 11:604913. doi:10.3389/fimmu.2020.604913

37. Talker SC, Koinig HC, Stadler M, Graage R, Klingler E, Ladinig A, et al. Magnitude and kinetics of multifunctional CD4+ and CD8β+ T cells in pigs infected with swine influenza A virus. Vet Res. (2015) 46:52. doi:10.1186/s13567-015-0182-3

38. Brown DM, Lee S, Garcia-Hernandez M de la L, Swain SL. Multifunctional CD4 Cells Expressing Gamma Interferon and Perforin Mediate Protection against Lethal Influenza Virus Infection. J Virol. (2012) 86:6792–803. doi:10.1128/JVI.07172-11

39. McMaster SR, Wilson JJ, Wang H, Kohlmeier JE. Airway-Resident Memory CD8 T Cells Provide Antigen-Specific Protection against Respiratory Virus Challenge through Rapid IFN-γ Production. J Immunol (2015) 195:203–9. doi:10.4049/jimmunol.1402975

40. Palomino-Segura M, Latino I, Farsakoglu Y, Gonzalez SF. Early production of IL-17A by γδ T cells in the trachea promotes viral clearance during influenza infection in mice. Eur J Immunol. (2020) 50:97–109. doi:10.1002/eji.201948157

41. Nazki S, Khatun A, Jeong CG, Mattoo SUS, Gu S, Lee SI, et al. Evaluation of local and systemic immune responses in pigs experimentally challenged with porcine reproductive and respiratory syndrome virus. Vet Res. (2020) 51:66. doi:10.1186/s13567-020-00789-7

42. Sedlak C, Patzl M, Saalmüller A, Gerner W. CD2 and CD8α define porcine γδ T cells with distinct cytokine production profiles. Dev Comp Immunol. (2014) 45:97–106. doi:10.1016/j.dci.2014.02.008

43. Pol JM, van Leengoed LA, Stockhofe N, Kok G, Wensvoort G. Dual infections of PRRSV/influenza or PRRSV/Actinobacillus pleuropneumoniae in the respiratory tract. Vet Microbiol. (1997) 55:259–64. doi:10.1016/s0378-1135(96)01323-5

44. Pomorska-Mól M, Podgórska K, Czyżewska-Dors E, Turlewicz-Podbielska H, Gogulski M, Włodarek J, Łukomska A. Kinetics of single and dual simultaneous infection of pigs with swine influenza A virus and porcine reproductive and respiratory syndrome virus. J Vet Intern Med. (2020) 34(5):1903–13. doi:10.1111/jvim.15832

45. Van Reeth K, Nauwynck H, Pensaert M. Clinical effects of experimental dual infections with porcine reproductive and respiratory syndrome virus followed by swine influenza virus in conventional and colostrum-deprived pigs. J Vet Med B Infect Dis Vet Public Health. (2001) 48:283–92. doi:10.1046/j.1439-0450.2001.00438.x

46. Kitikoon P, Vincent AL, Jones KR, Nilubol D, Yu S, Janke BH, et al. Vaccine efficacy and immune response to swine influenza virus challenge in pigs infected with porcine reproductive and respiratory syndrome virus at the time of SIV vaccination. Vet Microbiol. (2009) 139:235–44. doi:10.1016/j.vetmic.2009.06.003

47. Thacker EL, Thacker BJ, Young TF, Halbur PG. Effect of vaccination on the potentiation of porcine reproductive and respiratory syndrome virus (PRRSV)-induced pneumonia by Mycoplasma hyopneumoniae. Vaccine. (2000) 18:1244–52. doi:10.1016/s0264-410x(99)00395-3

48. Suradhat S, Kesdangsakonwut S, Sada W, Buranapraditkun S, Wongsawang S, Thanawongnuwech R. Negative impact of porcine reproductive and respiratory syndrome virus infection on the efficacy of classical swine fever vaccine. Vaccine. (2006) 24:2634–42. doi:10.1016/j.vaccine.2005.12.010

49. Oh T, Park KH, Yang S, Jeong J, Kang I, Park C, et al. Evaluation of the efficacy of a trivalent vaccine mixture against a triple challenge with Mycoplasma hyopneumoniae, PCV2, and PRRSV and the efficacy comparison of the respective monovalent vaccines against a single challenge. BMC Vet Res. (2019) 15:342. doi:10.1186/s12917-019-2091-6

50. Jeong J, Kim S, Park C, Park KH, Kang I, Park SJ, et al. Commercial porcine reproductive and respiratory syndrome virus (PRRSV)-2 modified live virus vaccine against heterologous single and dual Korean PRRSV-1 and PRRSV-2 challenge. Vet Rec. (2018) 182:485. doi:10.1136/vr.104397

51. Sattler T, Pikalo J, Wodak E, Revilla-Fernández S, Steinrigl A, Bagó Z, et al. Efficacy of live attenuated porcine reproductive and respiratory syndrome virus 2 strains to protect pigs from challenge with a heterologous Vietnamese PRRSV 2 field strain. BMC Vet Res. (2018) 14:133. doi:10.1186/s12917-018-1451-y

52. Fleming DS, Miller LC, Tian Y, Li Y, Ma W, Sang Y. Impact of porcine arterivirus, influenza B, and their coinfection on antiviral response in the porcine lung. Pathogens. (2020) 9:934. doi:10.3390/pathogens9110934

53. Munoz FM, Cramer JP, Dekker CL, Dudley MZ, Graham BS, Gurwith M, et al. Vaccine-associated enhanced disease: Case definition and guidelines for data collection, analysis, and presentation of immunization safety data. Vaccine. (2021) 39:3053–66. doi:10.1016/j.vaccine.2021.01.055

54. Costers S, Vanhee M, Van Breedam W, Van Doorsselaere J, Geldhof M, Nauwynck HJ. GP4-specific neutralizing antibodies might be a driving force in PRRSV evolution. Virus Res. (2010) 154:104–13. doi: 10.1016/j.virusres.2010.08.026

55. Zhang Z, Zhou L, Ge X, Guo X, Han J, Yang H. Evolutionary analysis of six isolates of porcine reproductive and respiratory syndrome virus from a single pig farm: MLV-evolved and recombinant viruses. Infect Genet Evol. (2018) 66:111–19. doi: 10.1016/j.meegid.2018.09.024

56. Nedumpun T, Sirisereewan C, Thanmuan C, Techapongtada P, Puntarotairung R, Naraprasertkul S, et al. Induction of porcine reproductive and respiratory syndrome virus (PRRSV)-specific regulatory T lymphocytes (Treg) in the lungs and tracheobronchial lymph nodes of PRRSV-infected pigs. Vet Microbiol. (2018) 216:13–19. doi:10.1016/j.vetmic.2018.01.014

57. Renukaradhya GJ, Alekseev K, Jung K, Fang Y, Saif LJ. Porcine reproductive and respiratory syndrome virus-Induced immunosuppression exacerbates the inflammatory response to porcine respiratory coronavirus in pigs. Viral Immunol. (2010) 23:457–66. doi:10.1089/vim.2010.0051

58. Subramaniam S, Kwon B, Beura LK, Kuszynski CA, Pattnaik AK, Osorio FA. Porcine reproductive and respiratory syndrome virus non-structural protein 1 suppresses tumor necrosis factor-alpha promoter activation by inhibiting NF-κB and Sp1. Virology. (2010) 406:270–9. doi:10.1016/j.virol.2010.07.016

59. Ke H, Han M, Zhang Q, Rowland R, Kerrigan M, Yoo D. Type I interferon suppression-negative and host mRNA nuclear retention-negative mutation in nsp1β confers attenuation of porcine reproductive and respiratory syndrome virus in pigs. Virology. (2018) 517:177–87. doi:10.1016/j.virol.2018.01.016

60. Hartmann W, Brunn ML, Stetter N, Gagliani N, Muscate F, Stanelle-Bertram S, et al. Helminth Infections Suppress the Efficacy of Vaccination against Seasonal Influenza. Cell Rep. (2019) 29:2243–2256.e4. doi:10.1016/j.celrep.2019.10.051

61. Janke B. H. (2013). Clinicopathological features of Swine influenza. Curr Top Microbiol Immunol. 370:69–83. doi:10.1007/82_2013_308

62. Rajao DS, Anderson TK, Kitikoon P, Stratton J, Lewis NS, Vincent AL. Antigenic and genetic evolution of contemporary swine H1 influenza viruses in the United States. Virology. (2018) 518:45–54. doi:10.1016/j.virol.2018.02.006

63. Mancera Gracia JC, Pearce DS, Masic A, Balasch M. Influenza A Virus in Swine: Epidemiology, Challenges and Vaccination Strategies. Front Vet Sci. (2020) 7:647. doi:10.3389/fvets.2020.00647

64. Huang KY, Rijal P, Schimanski L, Powell TJ, Lin TY, McCauley JW, et al. Focused antibody response to influenza linked to antigenic drift. J Clin Invest. (2015) 125:2631–45. doi:10.1172/JCI81104

65. Lorbach JN, Nelson SW, Lauterbach SE, Nolting JM, Kenah E, McBride DS, et al. Influenza Vaccination of Swine Reduces Public Health Risk at the Swine-Human Interface. mSphere. (2021) 6:e0117020. doi:10.1128/mSphere.01170-20

