## Supplementary Figure S1 for "Simultaneous infection with porcine reproductive and respiratory syndrome and influenza viruses abrogates clinical protection induced by live attenuated porcine reproductive and respiratory syndrome vaccination"

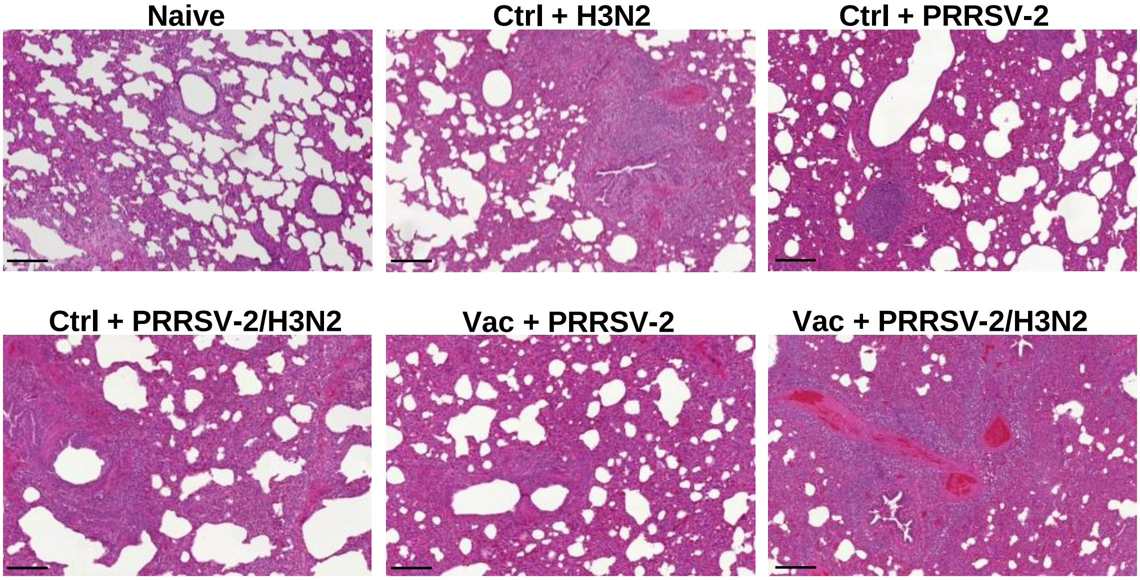


**Supplementary Figure 1. Histopathology of lungs.** Sections of cranial, cardiac and diaphragmatic lung lobes collected at 5 dpc were stained with H&E and microscopic lesions scored. Representative images of histologic samples from each group (n=6 per group) are shown (original magnification x 100; bar 100 μm).
