## Supplementary Figure S2 for "Simultaneous infection with porcine reproductive and respiratory syndrome and influenza viruses abrogates clinical protection induced by live attenuated porcine reproductive and respiratory syndrome vaccination"

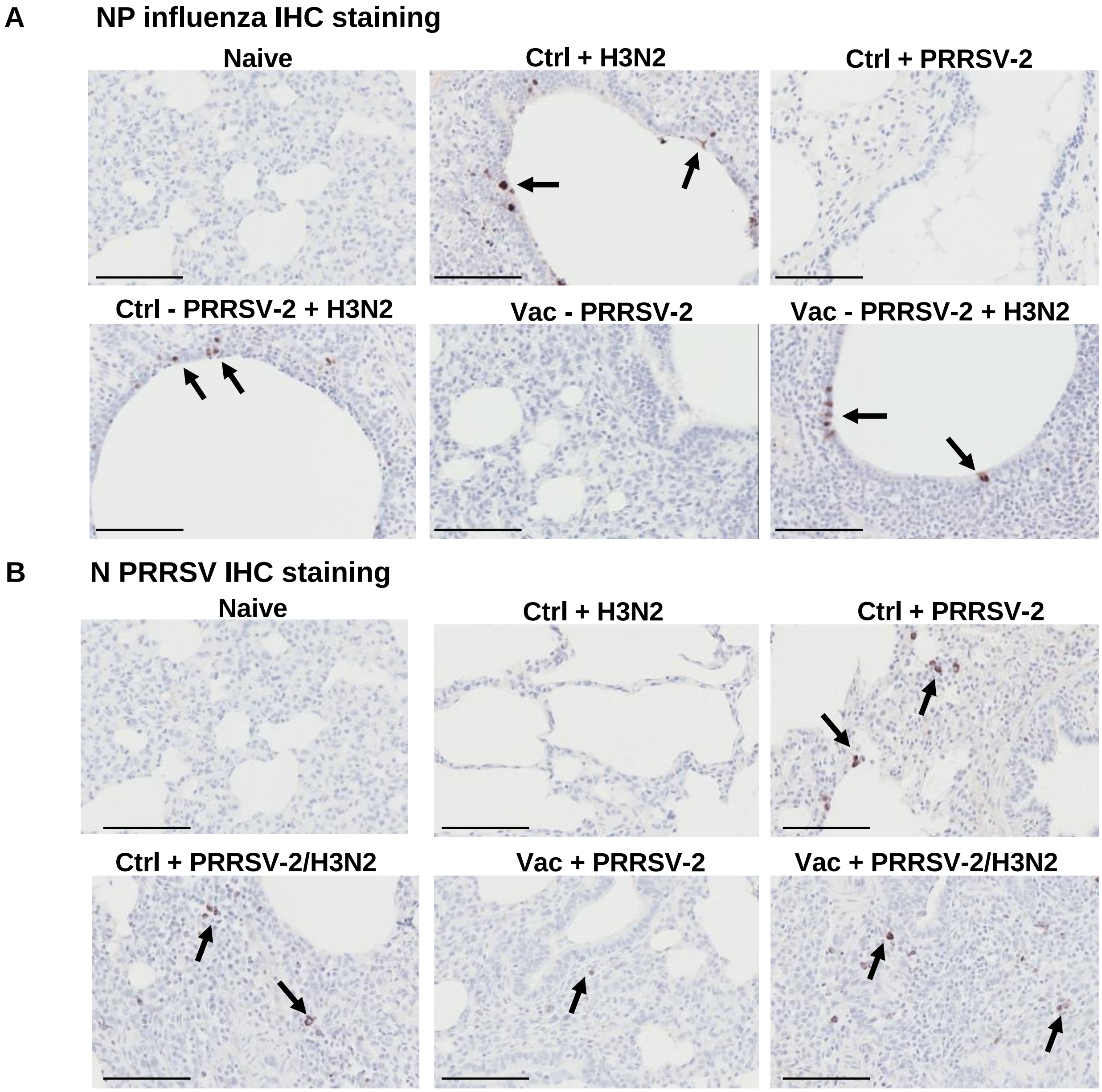


**Supplementary Figure 2. Immunohistochemical staining for virus detection.** Sections of of cranial, cardiac and diaphragmatic lung lobes collected at 5 dpc were stained for virus. **(A)** Lung sections stained using an anti-influenza NP mAb are shown and presence of NP-positive cells in bronchiolar epithelial cells are indicated by the black arrows. **(B)** Lung sections stained with an anti-PRRSV N mAb are shown and NP-positive cells in monocytes/macrophages are indicated by the black arrows. Immunohistochemical staining of representative lungs for each group (n=6 per group) are shown (original magnification x 400; scale bar 200 μm).
