## Supplementary Figure S3 for "Simultaneous infection with porcine reproductive and respiratory syndrome and influenza viruses abrogates clinical protection induced by live attenuated porcine reproductive and respiratory syndrome vaccination"

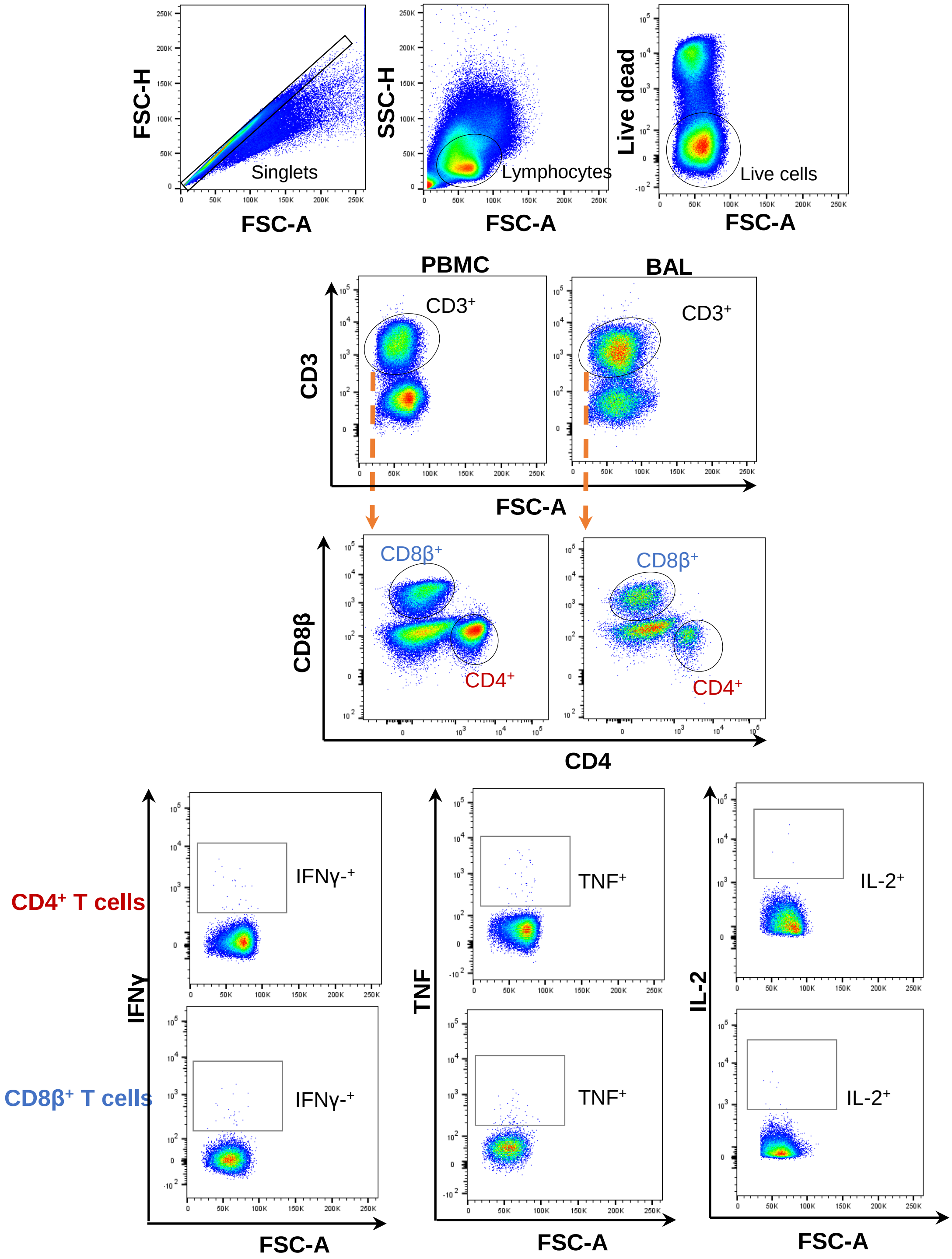


**Supplementary Figure 3. Intracellular cytokine staining (ICS) gating strategy.** Successive gates were applied to identify singlet cells, FSC/SSC defined lymphocytes, live, $CD3^+$ and subsequently $CD4^+$ or $CD8\beta^+$ T cells in the PBMC and BALF cells (upper and middle quadrant). IFN-γ-, TNF- and IL-2-producing cells within $CD4^+$ and $CD8\beta^+$ T cells are indicated in the lower quadrants.
