## Supplementary Figure S4 for "Simultaneous infection with porcine reproductive and respiratory syndrome and influenza viruses abrogates clinical protection induced by live attenuated porcine reproductive and respiratory syndrome vaccination"

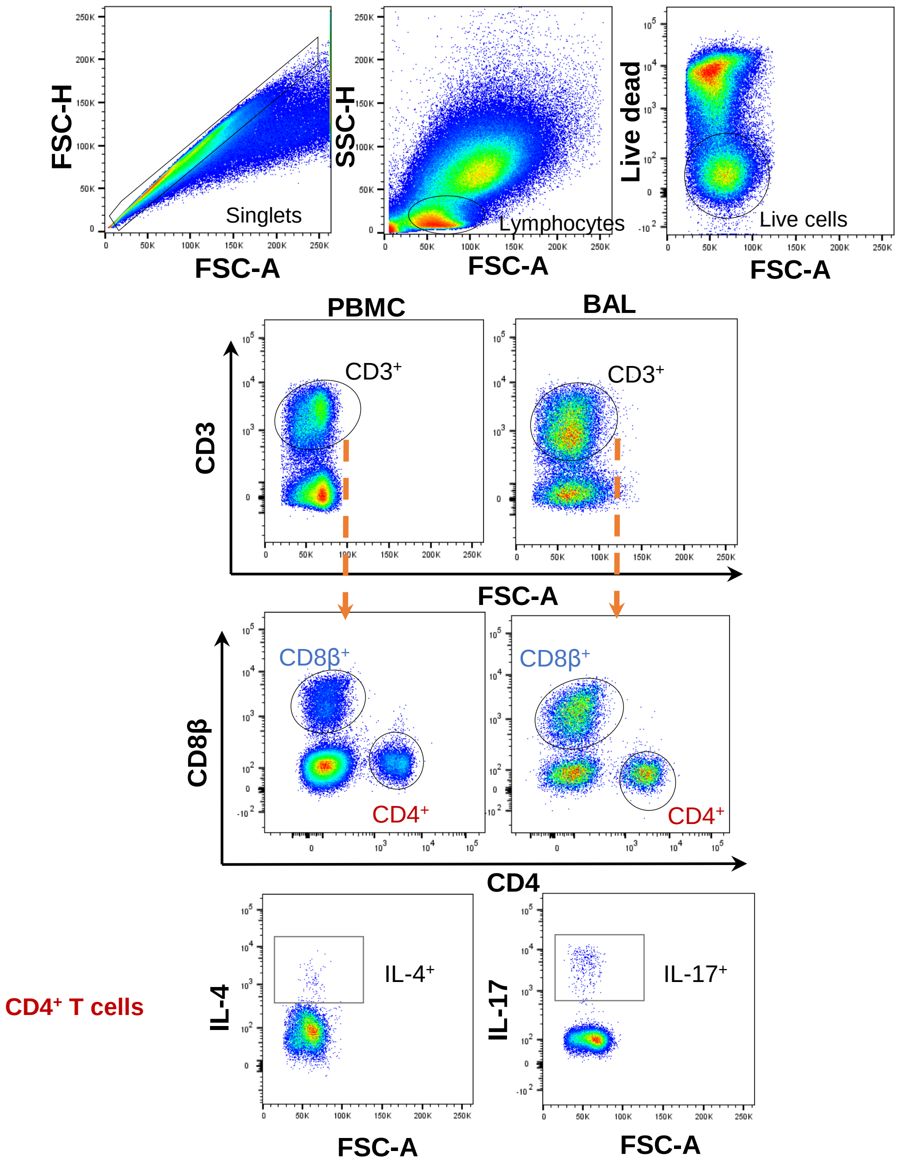


**Supplementary Figure 4. Intracellular cytokine staining (ICS) gating strategy.** Successive gates were applied to identify singlet cells, FSC/SSC defined lymphocytes, live, $CD3^{+}$ and subsequently $CD4^{+}$ or $CD8\beta^{+}$ T cells in the PBMC and BAL cells. IL-4- and IL-17- producing cells within $CD4^{+}$ T cells are indicated in the lower quadrants.
