## Supplementary Figure S5 for "Simultaneous infection with porcine reproductive and respiratory syndrome and influenza viruses abrogates clinical protection induced by live attenuated porcine reproductive and respiratory syndrome vaccination"

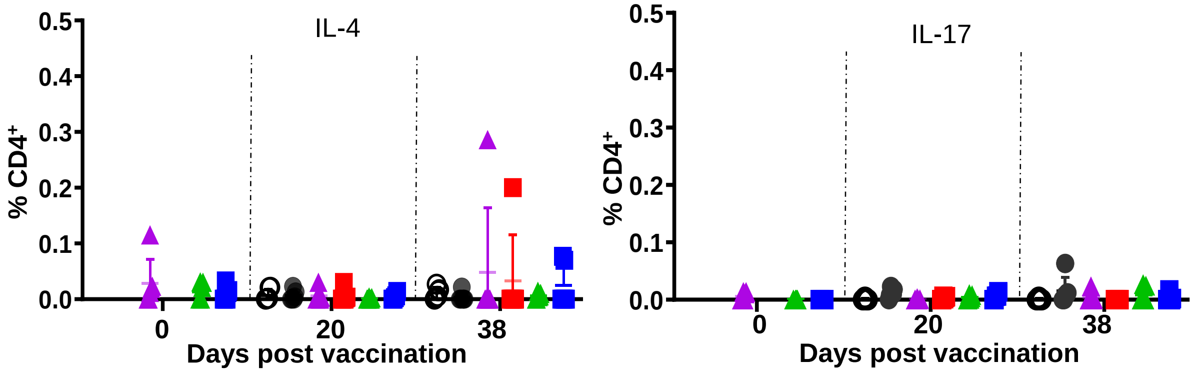


**Supplementary Figure 5. T cell responses against PRRSV-2.** PBMC cells were restimulated *in vitro* with PRRSV-2 or cultured with medium as previously described in **Figure 4**. Frequency of IL-4- and IL-17- secreting cells within the $CD4^+$ and $CD8\beta^+$ T cells are shown. The corrected frequencies (percentage of cytokine-producing cells subtracted with medium only) of each individual pigs and the mean + SD are displayed (n=5-6 per group).
