## Supplementary Figure S6 for "Simultaneous infection with porcine reproductive and respiratory syndrome and influenza viruses abrogates clinical protection induced by live attenuated porcine reproductive and respiratory syndrome vaccination"

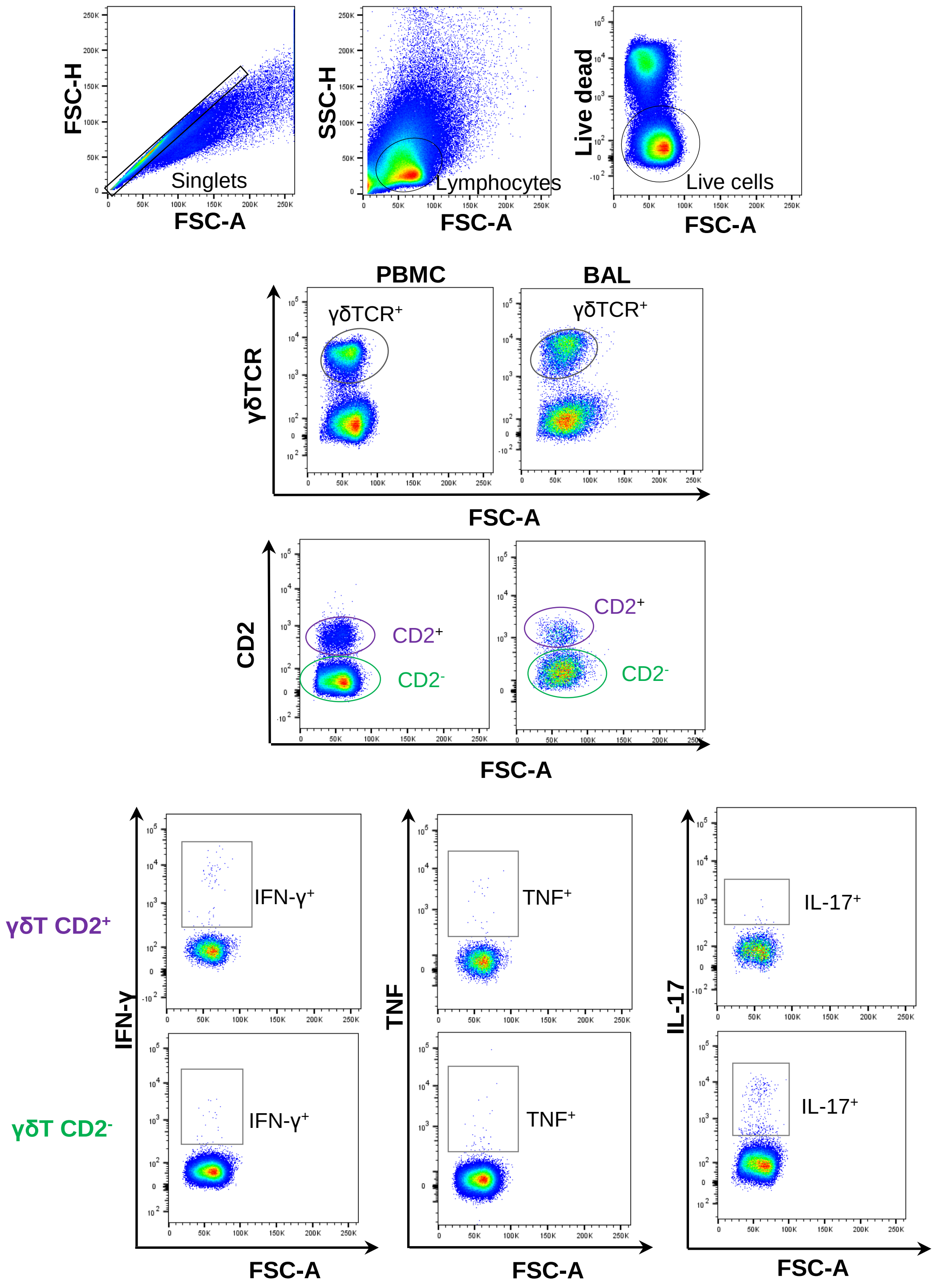


**Supplementary Figure 6. Intracellular cytokine staining (ICS) gating strategy.** Successive gates were applied to identify singlet cells, FSC/SSC defined lymphocytes, live, γδTCR$^+$ and subsequently CD2$^+$ or CD2$^-$ cells in the PBMC and BAL cells. IFN-γ-, TNF- and IL-17- producing cells within CD2$^+$ and CD2$^-$ γδT cells are indicated in the lower quadrants.
