## Supplementary Table S1 for "Simultaneous infection with porcine reproductive and respiratory syndrome and influenza viruses abrogates clinical protection induced by live attenuated porcine reproductive and respiratory syndrome vaccination"

**Supplementary Table 1. Scoring index of the clinical signs**

| Signs | Score |
|---|---|
| Temperature | 0-5 |
| Inappetence | 0-6 |
| Recumbancy | 0-6 |
| Skin discoloration | 0-3 |
| Respiratory changes | 0-6 |
| Nasal discharge | 0-2 |
| Eyes/conjunctiva | 0-1 |
| Body condition | 0-2 |
