## Supplementary Table S2 for "Simultaneous infection with porcine reproductive and respiratory syndrome and influenza viruses abrogates clinical protection induced by live attenuated porcine reproductive and respiratory syndrome vaccination"

**Supplementary Table 2. Antibodies used in flow cytometry**

| Panel | Antibody | Clone | Isotype | Specie | Dilution | Supplier |
|---|---|---|---|---|---|---|
| 1 | Anti-porcine CD3-PE | BB23-8E6-8C | IgG2a | Mouse | 1 : 200 | BD Biosciences |
| | Anti-porcine CD4-PerCP-Cy5.5 | 74-12-4 | IgG2b | Mouse | 1 : 100 | BD Biosciences |
| | Anti-porcine CD8β-FITC | PPT23 | IgG1 | Mouse | 1 : 400 | Bio-rad |
| | Anti-porcine IFN-γ-AF647 | P2G10 | IgG1 | Mouse | 1 : 800 | BD Biosciences |
| | Anti-human TNF-α-BV421 | Mab11 | IgG1 | Mouse | 1 : 100 | BioLegend |
| | Anti-porcine IL-2 | A150D3F1 | IgG2a | Mouse | 1 : 500 | Thermo Fisher Scientific |
| | Anti-mouse IgG2a-PE-Cy7 | m2a-15F8 | IgG1 | Rat | 1 : 400 | Thermo Fisher Scientific |
| 2 | Anti-porcine CD3-AF647 | BB23-8E6-8C | IgG2a | Mouse | 1 : 200 | BD Biosciences |
| | Anti-porcine CD4-PerCP-Cy5.5 | 74-12-4 | IgG2b | Mouse | 1 : 100 | BD Biosciences |
| | Anti-porcine CD8β-FITC | PPT23 | IgG21 | Mouse | 1 : 400 | Bio-rad |
| | Anti-human IL-17-PE | eBio64DEC17 | IgG1 | Mouse | 1 : 40 | Thermo Fisher Scientific |
| | Anti-human IL-4-BV421 | MP4-25D2 | IgG1 | Mouse | 1 : 40 | BioLegend |
| 3 | Anti-porcine CD2 | MSA4 | IgG2a | Mouse | 1 : 800 | Kingfisher Biotech |
| | Anti-porcine TCR1 Delta Chain | PGBL22A | IgG1 | Mouse | 1 : 800 | Kingfisher Biotech |
| | Anti-Mouse IgG2a-PE-Cy7 | m2a-15F8 | IgG1 | Rat | 1 : 400 | Thermo Fisher Scientific |
| | Anti-mouse IgG1- PerCP-Cy5.5 | RMG1-1 | IgG | Rat | 1 : 200 | BioLegend |
| | Anti-porcine IFN-γ-PE | P2G10 | IgG1 | Mouse | 1 : 800 | BD Biosciences |
| | Anti-human TNF-α-BV421 | Mab11 | IgG1 | Mouse | 1 : 100 | BioLegend |
| | Anti-porcine IL-17-FITC | eBio64DEC17 | IgG1 | Mouse | 1 : 40 | Thermo Fisher Scientific |
