## Supplementary Table S3 for "Simultaneous infection with porcine reproductive and respiratory syndrome and influenza viruses abrogates clinical protection induced by live attenuated porcine reproductive and respiratory syndrome vaccination"

### Supplementary Table 3. Body temperature recordings (°C) after the challenge

| | | Naïve | | | | | | Ctrl + H3N2 | | | | | | Ctrl + PRRSV-2 | | | | | | Ctrl + PRRSV-2/H3N2 | | | | | | Vac + PRRSV-2 | | | | | | Vac + PRRSV-2/H3N2 | | | | | |
|---|---|---|---|---|---|---|---|---|---|---|---|---|---|---|---|---|---|---|---|---|---|---|---|---|---|---|---|---|---|---|---|---|---|---|---|---|---|
| | Pig # | 1 | 2 | 3 | 4 | 5 | 6 | 7 | 8 | 9 | 10 | 11 | 12 | 13 | 14 | 15 | 16 | 17 | 18 | 19 | 20 | 21 | 22 | 23 | 24 | 25 | 26 | 27 | 28 | 29 | 30 | 31 | 32 | 33 | 34 | 35 | 36 |
| DPC[a] | -1 | 39.7 | 39.1 | 38.8 | 39.7 | 39.5 | 39.3 | 39.3 | 39.5 | 39.3 | 39.3 | 39.4 | 39.4 | 39.2 | 38.8 | 39.8 | 39.5 | 38.8 | 38.8 | 38.0 | 39.1 | 38.7 | 39.2 | 38.7 | 38.8 | 39.1 | 39.7 | 39.3 | 38.8 | 38.8 | 38.7 | 38.3 | 38.6 | 39.1 | 38.9 | 38.3 | 38.8 |
| | 0 | 38.9 | 39.6 | 39.2 | 38.6 | 39.2 | 39.0 | 38.5 | 38.3 | 38.7 | 38.9 | 39.0 | 38.8 | 39.3 | 39.2 | 39.6 | 39.5 | 38.8 | 39.6 | 38.8 | 39.2 | 39.2 | 38.7 | 38.9 | 39.3 | 38.6 | 38.3 | 38.3 | 38.4 | 38.4 | 38.4 | 38.9 | 39.2 | 38.6 | 39.0 | 38.6 | 38.5 |
| | 1 | 39.7 | 39.8 | 39.2 | 39.5 | 39.8 | 39.7 | 39.6 | 39.4 | 38.8 | 39.1 | 39.2 | 39.7 | 39.3 | 39.2 | 39.8 | 39.3 | 39.3 | 39.6 | 39.1 | 39.1 | 39.3 | 39.2 | 38.9 | 39.0 | 39.3 | 39.7 | 39.6 | 39.3 | 39.4 | 39.3 | 39.7 | 39.5 | **40.0** | 39.1 | 39.3 | 39.6 |
| | 2 | 39.7 | 39.5 | 39.1 | 39.5 | 39.8 | 39.4 | 39.3 | 38.9 | 38.4 | 39.6 | 39.2 | **40.0** | 38.1 | 39 | 39.5 | 39.2 | 38.9 | 39.3 | 38.8 | 38.6 | 39.0 | 39.0 | 39.0 | 39.6 | 39.4 | 39.7 | 39.7 | 39.6 | 39.9 | 39.4 | **40.3** | **40.1** | 39.3 | **40.9** | 39.3 | 39.5 |
| | 3 | 39.7 | 39.8 | 39.4 | 39.5 | 39.7 | 39.7 | 38.8 | 38.7 | 39.0 | 38.4 | 38.8 | 38.6 | 39.8 | 39.6 | 39.3 | 39 | 38.8 | 39.7 | 38.9 | 39.7 | 39.5 | 39.8 | 39.7 | 35.9 | 38.9 | 39.5 | 39.5 | 39.4 | 39.7 | 39.0 | 39.2 | 39.1 | 39.5 | **40.6** | 39.5 | 39.3 |
| | 4 | 39.2 | 39.6 | 39.1 | 39.4 | 39.5 | 39.4 | 39.3 | 39.0 | 39.3 | 38.9 | 38.9 | 39.0 | **40.1** | 39.9 | **40.0** | 39.4 | 38.9 | 39.9 | 38.9 | **40.4** | 39.4 | 39.6 | 38.9 | 39.5 | 39.1 | 39.1 | 38.4 | 38.6 | 38.6 | 38.2 | 39.6 | 39.6 | 39.3 | **40.7** | 39.3 | 39.5 |

a: days post challenge
