## Supplementary Table S4 for "Simultaneous infection with porcine reproductive and respiratory syndrome and influenza viruses abrogates clinical protection induced by live attenuated porcine reproductive and respiratory syndrome vaccination"

**Supplementary Table 4. Percentage identity of amino acid sequences among PRRS MLV (Genbank AF066183.4) and PRRSV-2 16CB02 (Genbank MZ700336) strain proteins**

| Protein | Percentage identity |
|---|---|
| GP2 | 92% |
| GP3 | 86% |
| GP4 | 89% |
| GP5 | 87% |
| M | 96% |
